# Fc-Free Single Chain Antibody mRNA Therapy for Airway Infection of Multidrug-Resistant *Pseudomonas Aeruginosa*

**DOI:** 10.1101/2025.06.12.659416

**Authors:** Mao Kinoshita, Ken Kawaguchi, Nguyen BT. Le, Atsushi Kainuma, Teiji Sawa, Satoshi Uchida

## Abstract

With the growing threat of antimicrobial resistance (AMR), alternatives to antibiotics are urgently needed. In this study, we developed mRNA-based therapeutics encoding single-chain variable fragment (scFv) antibodies that target the type III secretion system of *Pseudomonas aeruginosa*, a key AMR pathogen. Delivered intravenously via lipid nanoparticles, the scFv mRNA effectively protected mice from airway infections by mitigating lung inflammation, reducing bacterial load, and improving survival. Notably, this approach proved effective in clinically relevant models involving immunocompromised mice infected with multidrug-resistant, *exoU*-positive (highly cytotoxic) clinical isolates. The study also explores the potential of Fc-free scFv formulations, leveraging mRNA technology to overcome their short half-life through sustained protein production. Importantly, Fc-free scFv antibodies migrated more efficiently from the bloodstream to the airway epithelium—the primary site of infection—than their Fc-conjugated counterparts, resulting in efficient therapeutic outcomes. Overall, mRNA-encoded Fc-free scFv antibodies represent a promising strategy for combating AMR infections.

## Introduction

The rapid proliferation of drug-resistant bacteria has emerged as a critical and escalating threat to global public health. In 2017, the World Health Organization identified twelve priority pathogens, among which *Acinetobacter baumannii*, *Pseudomonas aeruginosa* (*P. aeruginosa*), and *Enterobacteriaceae* were deemed most concerning^1^. A global analysis published in 2019 estimated that antimicrobial resistance (AMR) was directly responsible for 1.27 million deaths, with approximately 80% of these attributable to six bacterial species, including *Escherichia coli*, *Staphylococcus aureus*, and *P. aeruginosa*^2^. With the pace of antibiotic discovery stagnating, projections suggest that by 2050, deaths caused by multidrug-resistant infections could exceed those from cancer^3^. The report, released in September 2024, warns that over the next 25 years, antibiotic-resistant bacteria may directly cause more than 39 million deaths worldwide, with an additional 169 million deaths resulting from complications associated with such infections ^4^. Alarmingly, the development of novel antibiotics continues to lag behind the accelerating evolution of bacterial resistance, underscoring the urgent need for innovative alternative therapeutic strategies^5^.

*P. aeruginosa* poses a substantial threat among drug-resistant pathogens, particularly by causing opportunistic respiratory infections in ventilated patients, individuals with cystic fibrosis, and immunocompromised hosts^6,7^. Monoclonal antibodies represent a promising therapeutic strategy, offering pathogen-specific targeting with minimal off-target effects. Our research has identified the type III secretion system (T3SS) and its associated exotoxins as critical targets for antibody-based intervention, given their strong association with acute lung injury and increased mortality^8^. Notably, in preclinical models, antibodies directed against PcrV—a structural cap protein of the T3SS— markedly reduced lung pathology and improved survival outcomes by effectively blocking the translocation of all T3SS effector toxins into host cells^9,10^.

Despite ongoing efforts—including previous clinical trials involving the monoclonal anti-PcrV antibody^11,12^, which we helped develop—no immunization strategies targeting *P. aeruginosa* have reached clinical application^13^, even a quarter-century after the initial proof of concept was demonstrated in animal models^14^. One major barrier to the clinical translation of recombinant antibodies for infectious diseases is the high cost associated with their complex production, purification, and characterization. In this context, mRNA therapy offers a significant advantage through its cell-free, streamlined manufacturing process, which simplifies both production and purification^15,16^. Unlike recombinant antibodies, which exhibit diverse physicochemical properties depending on their sequences, mRNAs encoding different antibodies share similar characteristics, enabling standardized manufacturing processes. Following the success of mRNA-based COVID-19 vaccines, there has been growing interest in using mRNA to produce therapeutic proteins in situ within the patient’s body. This approach has shown promise in clinical trials for genome editing and protein replacement therapies^17–19^. The application of mRNA is now expanding into antibody-based therapies, driving animal and clinical studies^20–26^. Additional advantages of mRNA include the ability to produce antibodies with patient-specific post-translational modifications and the ease of adapting the platform to antibody variants such as fragments and bispecifics.

In this context, we evaluated the therapeutic potential of mRNA encoding an anti-PcrV antibody in a murine model of *P. aeruginosa* infection. Leveraging the design flexibility of mRNA, we prepared two formats: scFvs with and without conjugation to the mouse IgG Fc domain (mFc). Notably, while recombinant Fc-free scFv antibodies typically exhibit a short circulation half-life on the order of tens of minutes, mRNA delivery can extend their retention in the body^27^. Using ionizable lipid nanoparticles (LNPs) for intravenous delivery, which induces protein expression in the liver, both

Fc-free and Fc-conjugated scFv mRNAs significantly improved survival following bacterial challenge. Notably, this approach proved effective in a clinically relevant model involving immunocompromised mice infected with multidrug-resistant, highly cytotoxic clinical isolates of *P. aeruginosa*. Structural predictions and supporting studies indicate that mutations in these isolates, as well as other reported mutations, do not impair antibody recognition^28–30^, underscoring the robustness of our approach. Furthermore, our study revealed a key advantage of the Fc-free formulation over the Fc-conjugated version in terms of biodistribution. While current mRNA-based antibody therapies typically utilize Fc-conjugated formats^20,21,23,26,31^, we found that Fc-free scFv migrates from the bloodstream to the alveolar space, the primary site of *P. aeruginosa* infection more efficiently than Fc-conjugated scFv, resulting in efficient therapeutic outcomes.

**Figure 1.**
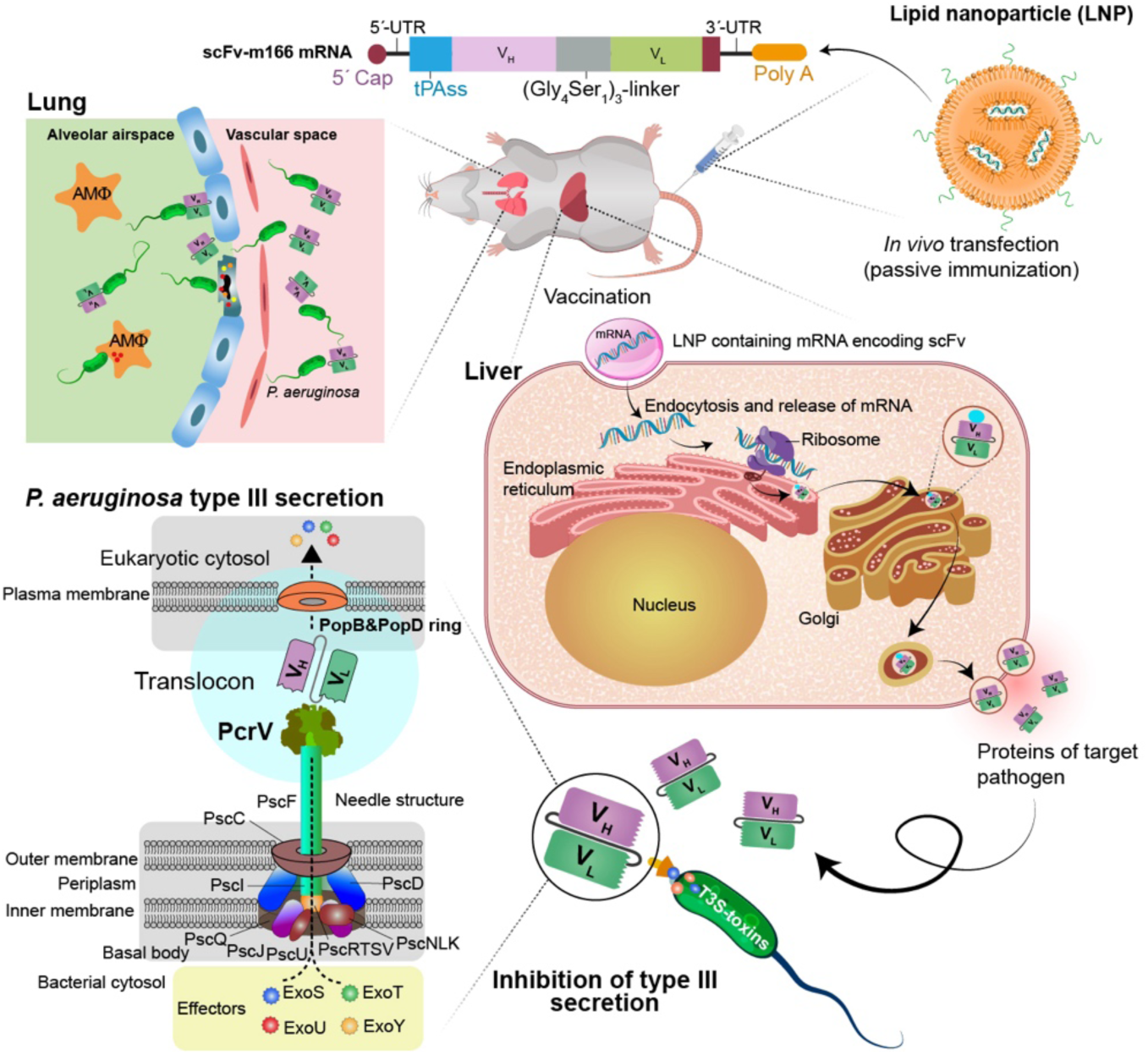
Schematic overview of this study. Intravenous administration of LNP encapsulating scFv mRNA targeting *P. aeruginosa* PcrV inhibits type III secretion system-mediated virulence, thereby protecting mice from bacterial challenge.

## Results

### Design of anti-PcrV scFv mRNA

We constructed mRNA encoding an anti-PcrV scFv antibody (scFv-m166) based on the genetic information of the murine monoclonal anti-PcrV IgG (mAb166)^32^ and added a tissue plasminogen activator signal sequence (tPAss) at the amino-terminus to facilitate secretion (**Fig. 2A-a**). To evaluate the influence of the Fc domain, we also prepared mRNA encoding tPAss-scFv conjugated with the mouse hinge region and the CH2 and CH3 Fc-domains at the carboxy-terminus (scFv-m166-mFc, **Fig. 2A-b**). As negative controls, we used mRNA encoding either luciferase or an irrelevant scFv targeting hen egg lysozyme (scFv-control, **Fig. 2A-c**)^33^. All four mRNA constructs were encapsulated in LNPs prior to injection into mice. The LNPs exhibited hydrodynamic diameters ranging from 100 to 130 nm with narrow polydispersity indexes below 0.15 in dynamic light scattering measurement and demonstrated high mRNA encapsulation efficiency above 90% (**Table S1**).

**Figure 2.**
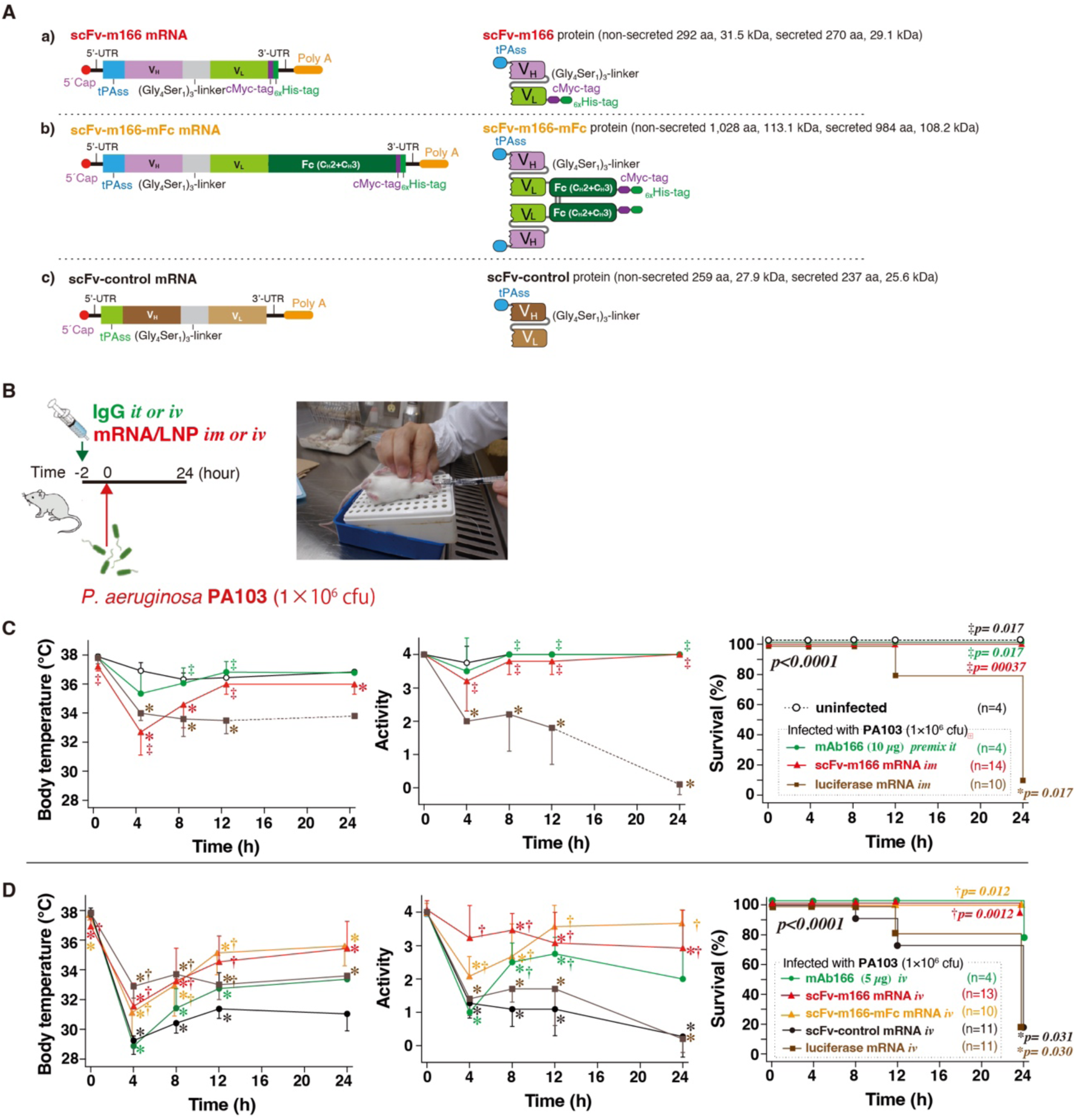
Design of anti-PcrV scFv mRNA constructs and prophylactic efficacy following bacterial challenge. **A.** Schematic representation of mRNA constructs: **a)** Fc-free **scFv-m166. b)** Fc-conjugated **scFv-m166-mFc** mRNA**. c)** Fc-free **scFv-control mRNA**. **B.** Experimental protocol for the prophylactic model. Bacteria were administered intratracheally (right). **C–D.** Post-challenge monitoring of body temperature (left), physical activity (center), and survival (right). **C.** Intramuscular injection. **D.** Intravenous injection. m166 IgG 10 µg premix it: Mice received *P. aeruginosa* pre-mixed with 10 µg anti-PcrV IgG protein (mAb166) intratracheally (it). mAb166 IgG iv: Mice received 5 µg mAb166 protein intravenously. \**P* < 0.05 vs. uninfected control; †*P* < 0.05 vs. intravenous scFv-control mRNA; ‡*P* < 0.05 vs. intramuscular luciferase mRNA.

### Prophylactic administration of anti-PcrV scFv antibody mRNA

We first investigated the therapeutic potential and mode of action of scFv-m166 and scFv-m166-mFc mRNA in a prophylactic setting. In this model, mice received mRNA LNPs either intramuscularly or intravenously two hours prior to bacterial challenge. As a control, anti-PcrV IgG (mAb166) protein was administered intravenously. Based on a previous preclinical study of mRNA-based antibody therapy for infectious disease, we used a dose of 10 µg/mouse for mRNA and 5 or 10 µg/mouse for mAb166 IgG protein throughout this study^21^. For the challenge, mice were intratracheally instilled with a lethal dose (1×10⁶ CFU/mouse) of *P. aeruginosa* PA103 (**Fig. 2B**). Previous studies have shown that, without treatment, most mice succumb within 24 hours due to acute lung injury caused by type III-secreted bacterial toxins^8,34^. Therefore, the observation period for this experiment was set to 24 hours.

In the intramuscular mRNA treatment, scFv-m166 mRNA conferred 100% survival at 24 hours post-challenge, whereas the survival rate in the control group receiving *luciferase* mRNA was below 10% (**Fig. 2C**). Consistent with this, scFv-m166 mRNA mitigated the drop in body temperature and activity levels following infection, indicating that the antibody produced from the mRNA effectively protected against *P. aeruginosa* lung infection. As a positive control, the “mAb166 IgG premix it” group received the lethal bacterial dose pre-mixed with recombinant antibody via intratracheal (it) instillation. All mice in this group survived the 24-hour period, confirming the neutralizing activity of the antibody.

mRNA-based antibody therapy also demonstrated efficacy following intravenous administration. Intravenous injection of mRNA LNPs led to predominant protein expression in the liver, particularly in hepatocytes (**Fig. S1**). In this setting, both scFv-m166 and scFv-m166-mFc mRNAs significantly improved mouse survival and preserved body temperature and activity levels compared to control groups receiving *luciferase* or scFv-control mRNA (**Fig. 2D**).

We further explored the mode of action. Mice injected with *luciferase* or scFv-control mRNA showed significant increases in lung weight, neutrophil myeloperoxidase (MPO) activity, and bacterial load in the lungs compared to uninfected controls at 24 hours post-challenge (**Figs. 3A-C**). The increase in lung weight indicates edema, while the elevated MPO activity reflects inflammatory responses. Notably, both intramuscular and intravenous administration of scFv-m166 or scFv-m166-mFc mRNA effectively alleviated these pathological changes and reduced bacterial loads in the lungs. Consistent with these findings, mRNA treatment minimized the expression of interleukin-6 (IL-6) and tumor necrosis factor-α (TNF-α) in lung homogenates to levels comparable to those in uninfected controls, whereas control mRNA groups showed elevated cytokine levels (**Figs. 3D, E**).

**Figure 3.**
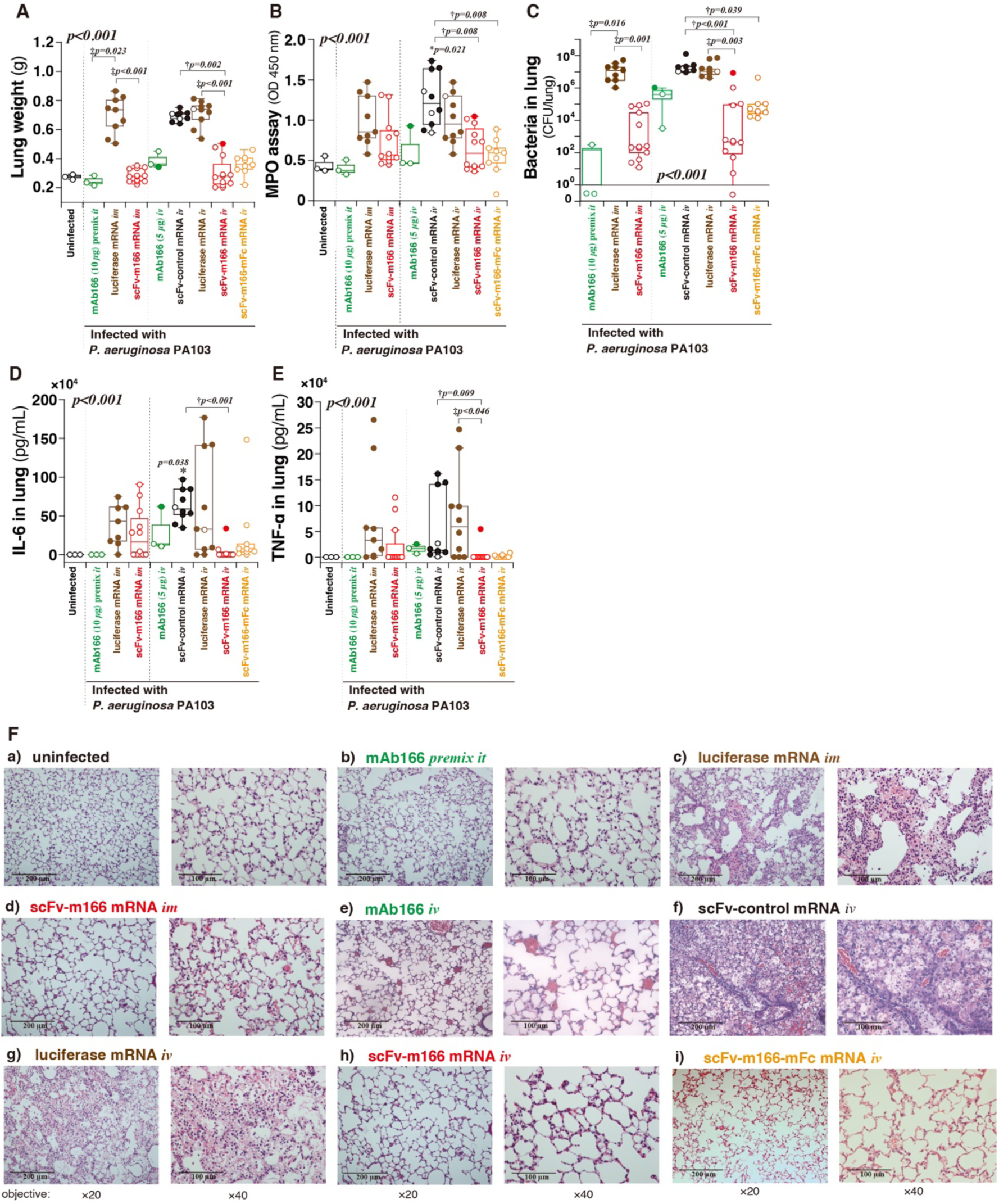
Mode of action of scFv mRNA in the prophylactic model. Mice were challenged with *P. aeruginosa* two hours after mRNA administration. Parameters assessed 24 hours post-challenge: **A.** Wet lung weight, reflecting pulmonary edema; **B.** Myeloperoxidase (MPO) activity; **C.** Bacterial burden (colony-forming units (CFU) per lung); **D.** Interleukin-6 (IL-6) concentration; **E.** Tumor necrosis factor-alpha (TNF-α) concentration in lung homogenates. Data are presented as box plots showing the interquartile range (box), median (centerline), and minimum and maximum values (whiskers). \**P* < 0.05 vs. uninfected control; †*P* < 0.05 vs. intravenous scFv-control mRNA; ‡*P* < 0.05 vs. luciferase mRNA. **F.** Representative hematoxylin and eosin (H&E) staining of lung sections. Left: ×20 objective magnification (scale bar: 200 µm); right: ×40 magnification (scale bar: 100 µm).

Histological analysis at 24 hours post-challenge further confirmed the therapeutic effects (**Fig. 3F**). Lungs from mice treated with *luciferase* or scFv-control mRNA exhibited signs of acute lung injury, including marked cellular infiltration, alveolar hemorrhage, and atelectasis. In contrast, treatment with scFv-m166 or scFv-m166-mFc mRNA—via either intramuscular or intravenous routes—alleviated these pathological changes, rendering lung histology nearly indistinguishable from that of uninfected mice.

These experiments using a prophylactic model of *P. aeruginosa* infection demonstrated the effectiveness of mRNA-encoding anti-PcrV Fc-free and Fc-conjugated antibodies in reducing bacterial load and mitigating lung inflammation, ultimately minimizing mortality (**Figs. 2 and 3**). While both intramuscular and intravenous routes were effective for delivering mRNA LNPs, intravenous injection may be more suitable for future applications in larger animals and humans. In larger animals, the relative distribution volume of intramuscularly injected solutions compared to total body volume becomes smaller. In contrast, intravenous injection allows targeting of the liver—a large, accessible organ—and several clinical trials have demonstrated the feasibility of liver-specific delivery using LNPs in humans^17,18^. Therefore, we adopted the intravenous route for subsequent therapeutic experiments.

### Therapeutic administration of anti-PcrV scFv mRNA

Next, we evaluated the therapeutic potential of anti-PcrV scFv antibody mRNA by administering the mRNA 30 minutes after intratracheal challenge with *P. aeruginosa* PA103 (**Fig. 4A**). This model more closely simulates a clinical treatment scenario for *P. aeruginosa* infection compared to the prophylactic model described above (**Fig. 2, 3**). Mouse survival was monitored over the course of one week. In the control group treated with scFv-control mRNA, all mice died within 24 hours (**Fig. 4B**). In contrast, intravenous administration of mAb166 IgG protein improved survival in a dose-dependent manner, reaching up to 80% at a dose of 10 µg. Intriguingly, intravenous injection of scFv-m166 mRNA significantly improved mouse survival compared to the scFv-control group, achieving an 80% survival rate one week post-challenge (**Fig. 4C**). scFv-m166-mFc mRNA also significantly improved survival compared to scFv-control mRNA, although to a lesser extent than the Fc-free scFv-m166 mRNA, resulting in a 50% survival rate one week post-challenge.

**Figure 4.**
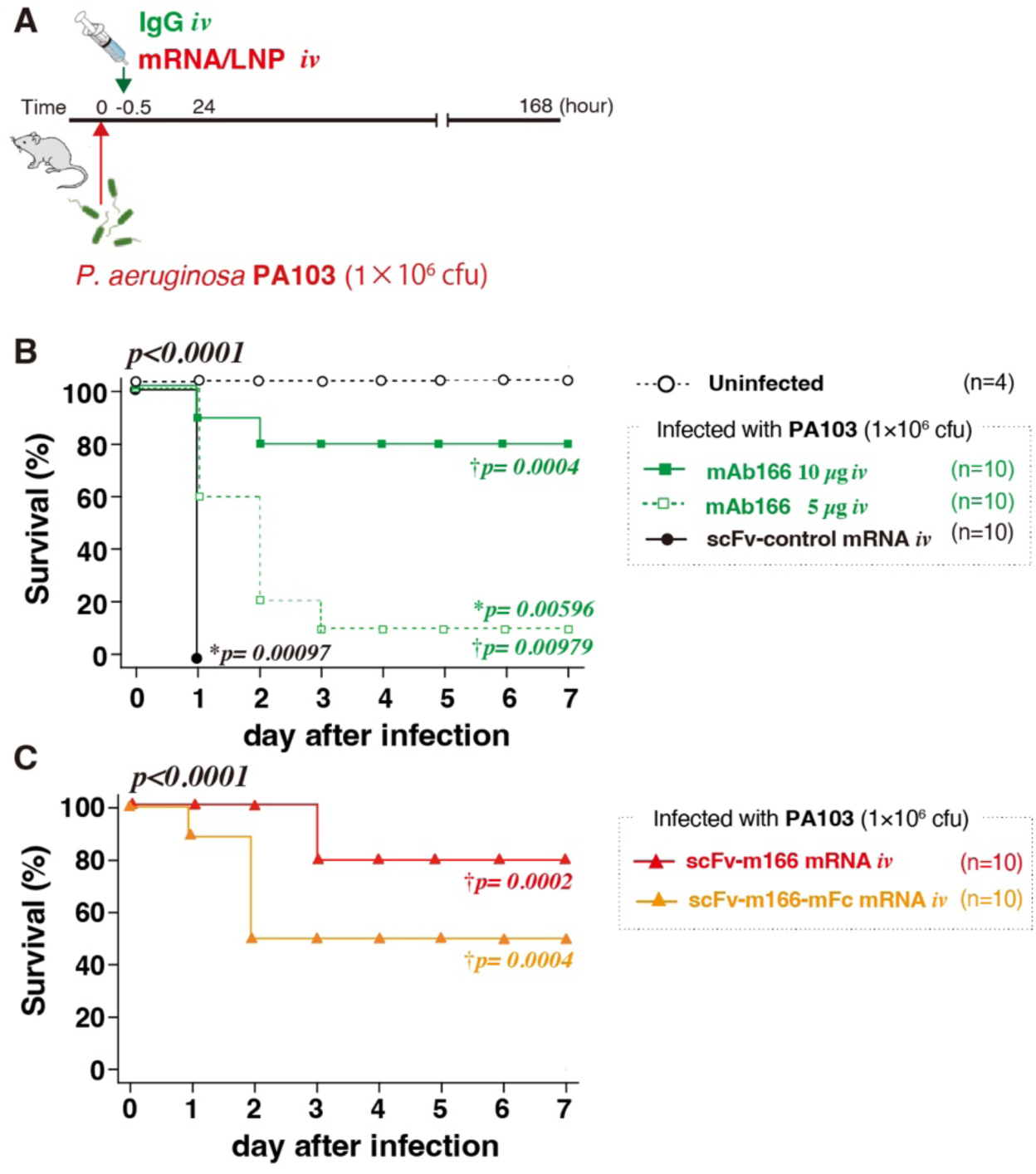
Therapeutic efficacy of mRNA treatment against *P. aeruginosa* infection. **A.** Experimental protocol. **B.** Survival of mice treated with 5 µg or 10 µg mAb166 IgG protein. **C.** Survival of mice treated with scFv-m166 or scFv-m166-mFc mRNA. \**P* < 0.05 vs. uninfected control; †*P* < 0.05 vs. scFv-control mRNA.

### Clinically isolated antimicrobial-resistant *P. aeruginosa* strains

Antibody therapy offers a promising solution to the growing clinical challenge of antimicrobial-resistant *P. aeruginosa*, as antibodies function through mechanisms distinct from those of antimicrobials. This motivated us to evaluate the feasibility of our approach using clinically isolated antimicrobial-resistant strains. To this end, we prepared 10 in-hospital isolates^35^, along with laboratory strains PAO1 and PA103. All 10 clinical isolates exhibited resistance to multiple antimicrobials, with varying susceptibility profiles. In contrast, the laboratory strain, PA103, used in the previous experiments (**Figs. 2-4**) was susceptible to all tested antimicrobials (**Fig. 5A**).

**Figure 5.**
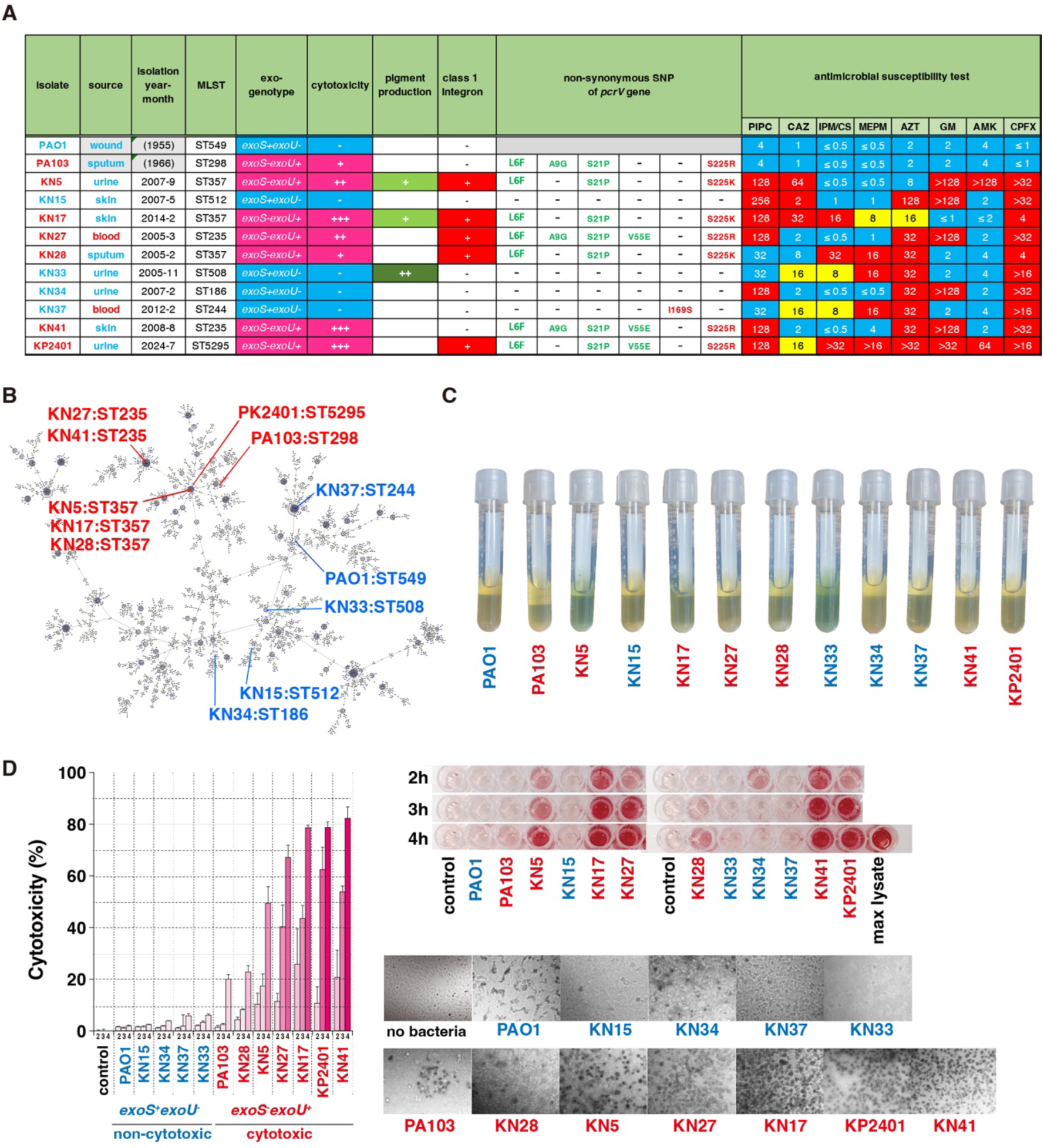
Characterization of various *P. aeruginosa* clinical isolates. **A.** Summary of 12 strains, including 2 laboratory strains (PAO1 and PA103) and 10 clinical isolates: multilocus sequence typing (MLST), exoenzyme genotypes, cytotoxicity, pigment production, non-synonymous SNPs in the *pcrV* gene, and antimicrobial susceptibility profiles. **B.** eBURST analysis of MLST data. A minimum spanning tree was constructed using PHYLOViZ data. **C.** Pigment production in tryptic soy broth. **D.** Cytotoxicity toward BEAS-2B bronchial epithelial cells. Right: quantitative data (mean ± SD of triplicates) 2 – 4 hours post-infection. Upper left: a representative image of the assay plate. Lower left: trypan blue staining at 4 hours, visualizing cell death. Non-cytotoxic strains (*exoS*⁺/*exoU*⁻): blue. Cytotoxic strains (*exoS*⁻/*exoU*⁺): red. MLST, multilocus sequence typing; PIPC, piperacillin; CAZ, ceftazidime; IPM/CS, imipenem/cilastatin; MEPM, meropenem; AZT, aztreonam; GM, gentamicin; AMK, amikacin; CPFX, ciprofloxacin.

Notably, *P. aeruginosa* possesses multiple virulence factors, and it is well-known that pathogenicity varies among isolates. In particular, T3SS-associated cytotoxicity, which correlates with lung injury and mortality, is linked to the *exoS*/*exoU* genotypes^34^. The clinical isolates differed in their *exoS*/*exoU* genotypes (**Figs. 5A and S2**). Multilocus sequence typing (MLST) analysis revealed that the 12 strains, including the 10 clinical isolates and two laboratory strains, were widely distributed across the eBURST phylogenetic tree, which includes 5,331 sequence types (STs) as reported in the PubMLST database at the time of analysis. Several isolates, including KN5, KN17, and KN33, exhibited high levels of pigment production in vitro (**Fig. 5C**), a trait often associated with increased virulence. The cytotoxicity of the 12 tested strains against lung epithelial cells is shown in **Fig. 5D**. Six clinical isolates exhibited enhanced cytotoxicity than the highly virulent laboratory strain PA103; all of these were of the *exoS*⁻/*exoU*⁺ genotype. In contrast, five strains, including PAO1, displayed lower cytotoxicity than PA103 and were all of the *exoS*⁺/*exoU*⁻ genotype.

To investigate whether mutations in the clinical isolates might affect antibody binding, we sequenced the *pcrV* genes from all 12 strains and compared the results with previously reported mutations known to disrupt recognition by the mAb166 blocking antibody^36^. Using the PcrV amino acid sequence of the PAO1 strain—serving as the reference in the *P. aeruginosa* whole-genome project—as a baseline, 1 to 5 amino acid substitutions were identified in 8 strains, including PA103. The locations of these mutations and their corresponding strains are illustrated in a molecular phylogenetic tree based on PcrV sequences (**Fig. 6A**). In total, seven amino acid substitutions were identified, including one previously unreported mutation within the m166 blocking epitope (I169S), along with two previously reported substitutions (S225K/R) ^36^. Bacterial proteins from the 12 *P. aeruginosa* strains cultured in Trypticase soy broth containing 10 mM nitrilotriacetic acid to induce T3SS expression were separated by SDS-PAGE and subjected to immunoblotting using the mAb166 IgG. Although considerable variation in PcrV expression levels was observed, all 12 strains displayed a band corresponding to PcrV at approximately 32.3 kDa (**Fig. 6B**). Structurally, PcrV is predicted to form a mushroom-shaped cap comprising a coiled-coil shaft—formed by central and C-terminal helical regions—and a globular domain of pentameric PcrV in complex with the scFv-m166 antibody (**Fig. 6C, D**)^28–30^. AlphaFold3-based structural modeling with Visual Molecular Dynamics (VMD) revealed that the mAb166 Fab and the engineered scFv-m166 predominantly bind to the globular domain at the periphery of the secretion pore and partly block the outlet of the secretion pore (**Figs. 6E, F, S3, and S4**). Further modeling using AlphFold3 indicated that the identified amino acid substitutions, including those within the blocking epitope, are located at positions spatially distant from the mAb166 binding interface and do not induce significant conformational changes to the mushroom-like cap structure of PcrV (**Fig. 6G, S4**). These findings support the structural robustness of scFv-m166 antibody binding across genetically diverse *P. aeruginosa* isolates.

**Figure 6.**
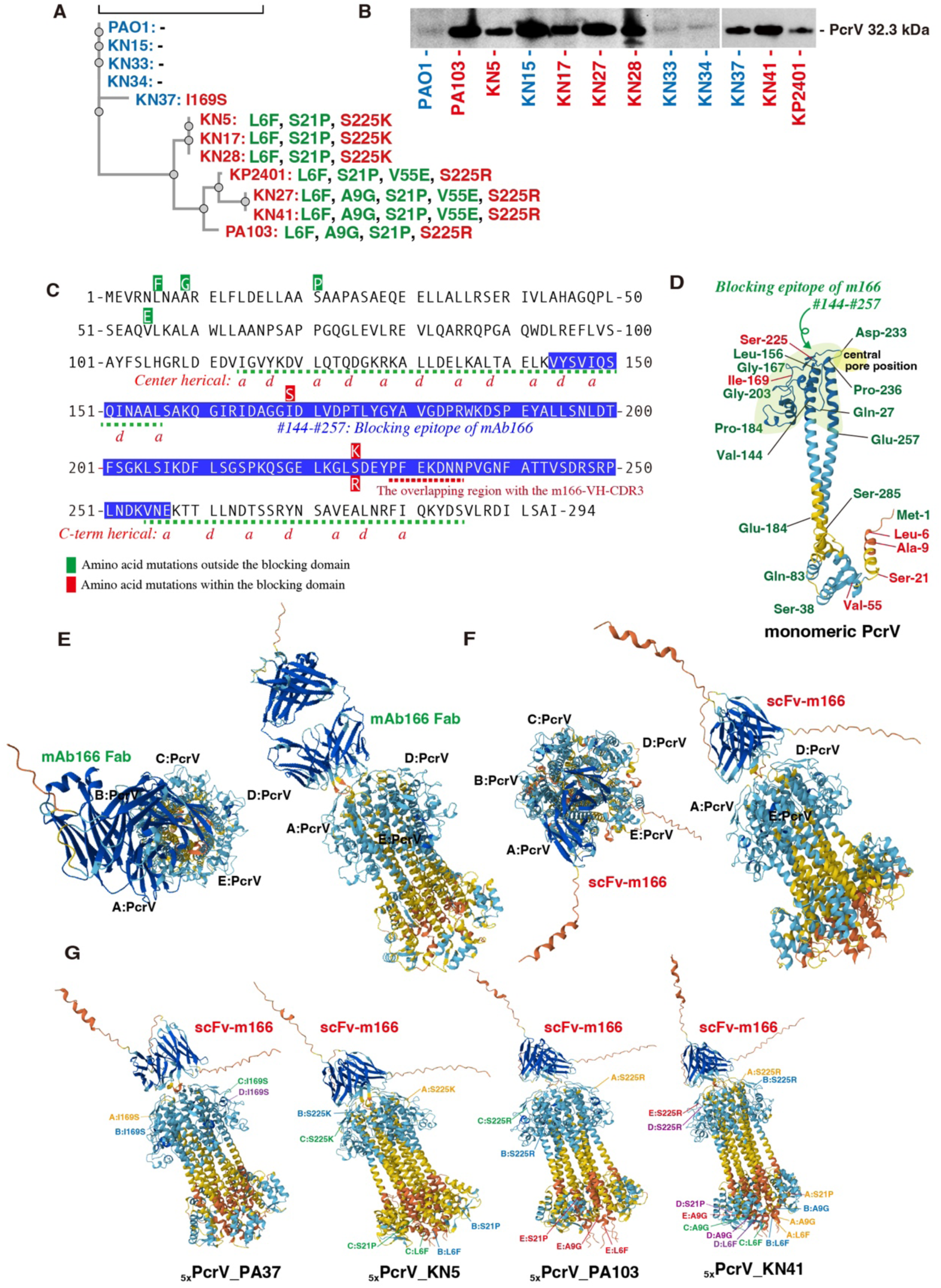
Phylogenetic analysis and 3D-structure prediction. **A.** Phylogenetic tree of PcrV associated non-synonymous SNPs among 12 strains. The anti-PcrV antibody (m166) recognizes blocking domain of PcrV (aa #144 – 257). Red: mutation within the domain; Green: mutations outside the domain. ***B.*** Immunoblot analysis of PcrV protein expression using anti-PcrV IgG (mAb166). **C.** PcrV sequence (PAO1), mutation sites, and blocking epitope recognized by mAb166 (blue). Green dotted lines: helical structure forming a coiled coil. Positions a and d among the seven amino acid residues (a to g) that constitute the helix are indicated. Red dotted line: overlap with m166-VH-CDR3 region. **D.** AlphaFold3 prediction of PcrV blocking domain. **E-G.** AlphaFold3 prediction of pentameric structure binding to the m166 Fab (**E**) and m166 scFv (**F, G**). **E, F.** Left: top views of PcrV bound to m166 Fab or scFv-m166. Right: side views of binding structure. **G.** The positions of non-synonymous SNPs.

### Therapeutic administration against various antimicrobial-resistant strains under immunocompromised conditions

Next, we evaluated the therapeutic potential of anti-PcrV scFv mRNA against *P. aeruginosa* infections using these clinical isolates. Since *P. aeruginosa* infections frequently occur in immunocompromised patients, we modeled this clinical scenario using immunocompromised mice. Immunosuppression was induced by intraperitoneal administration of cyclophosphamide, given two and four days prior to bacterial challenge (**Fig. 7A**). This regimen successfully induced leukopenia, as confirmed in **Fig. 7B**. To assess the broad efficacy of our approach, we conducted an initial screen using the 12 *P. aeruginosa* strains described earlier. In this setup, each group consisted of 12 mice, with each mouse infected with a different strain. Thirty minutes post-infection, mice received an intravenous injection of either scFv-control mRNA, scFv-m166 mRNA, scFv-m166-mFc mRNA, mAb166 IgG protein (10 µg), or colistin sulfate (125 µg). Survival was then monitored over 7 days.

**Figure 7.**
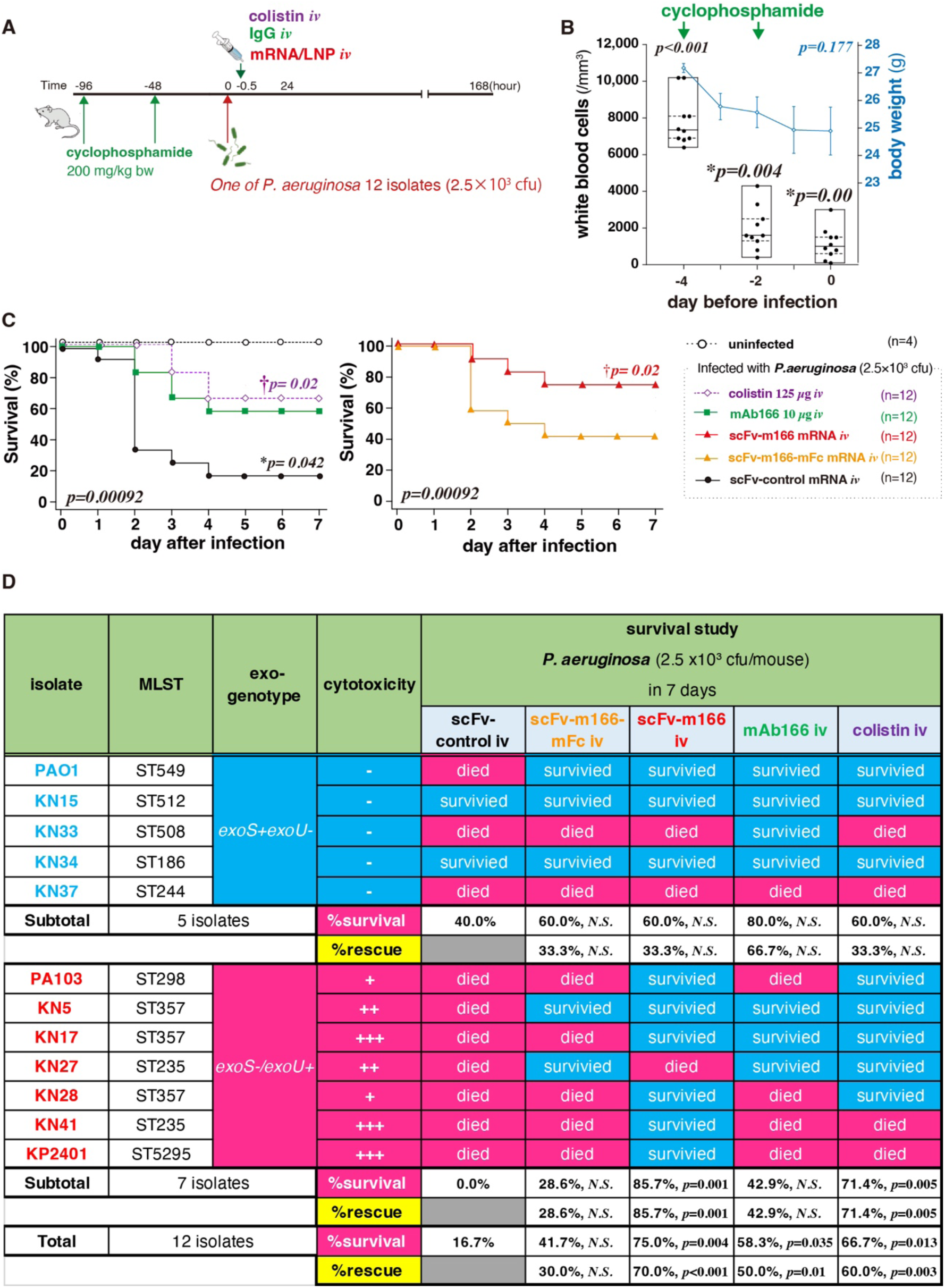
Therapeutic efficacy of treatment in immunocompromised mice infected with 12 clinical isolates of *P. aeruginosa*. **A.** Schematic of the experimental protocol. **B.** Effects of cyclophosphamide on peripheral blood leukocyte counts and body weight. Body weight changes are presented as mean ± SD. Leukocyte counts are displayed using five-number summaries (maximum, 75th percentile, median, 25th percentile, and minimum). *P* < 0.05 vs. untreated controls. **C.** Survival following bacterial challenge. The survival rate in the scFv-control group was significantly lower than that in the uninfected control group (*P* = 0.042). Both the colistin and scFv-m166 groups exhibited significantly improved survival compared with the scFv-control group (†*P* = 0.02). **D.** Summary of therapeutic outcomes. Strains were classified into low-virulence (*exoS*⁺/*exoU*⁻) and high-virulence (*exoS*⁻/*exoU*⁺) groups. %Rescue indicates the percentage of strains that were lethal in the scFv-control group but resulted in survival in the scFv-m166 group. Statistical analyses were performed using the Pearson Chi-square test. N.S.: Not significant (*P* ≥ 0.5).

Among the 12 mice treated with scFv-control mRNA, 10 died within 7 days, corresponding to a survival rate of 16.7% (**Figs. 7C and 7D**). In contrast, treatment with colistin sulfate (125 µg) resulted in 4 deaths (66.7% survival), and administration of mAb166 IgG protein (10 µg) led to 5 deaths (58.3% survival) over the same period. Notably, only 3 of the 12 mice treated with scFv-m166 mRNA died, yielding a significantly higher survival rate of 75.0% (*P* = 0.004). However, scFv-m166-mFc mRNA treatment resulted in 7 deaths (41.7% survival), indicating reduced efficacy. Among the 10 bacterial strains that caused death in the scFv-control mRNA group, scFv-m166 mRNA treatment rescued 7 mice, corresponding to a 70.0% rescue rate in otherwise lethal infections (*P* < 0.001) (**Fig. 7D**). Similarly, within the 7-day observation period, 5 mice (50%, *P* = 0.01) treated with mAb166 IgG protein and 6 mice (60.0%, *P* = 0.01) treated with colistin sulfate succumbed to infection. Importantly, focusing on infections caused by the 7 highly cytotoxic *exoU*^+^ clinical isolates, all mice treated with scFv-control mRNA died, whereas 6 of 7 mice (85.7%) treated with scFv-m166 mRNA survived (*P* = 0.001), highlighting the potent protective effect of scFv-m166 mRNA in severe infection settings.

To further evaluate the potential of our approach against highly cytotoxic and lethal *exoU+* clinical isolates, we conducted therapeutic experiments targeting the KN41 and KP2401 strains. These two strains exhibited the highest cytotoxicity toward BEAS-2B bronchial epithelial cells among the 12 strains tested (**Fig. 5D**). KN41 is resistant to piperacillin (PIPC), aztreonam (AZT), gentamicin (GM), and ciprofloxacin (CPFX) (**Fig. 5A**). KP2401, which is resistant to PIPC, imipenem/cilastatin (IPM/CS), meropenem (MEPM), AZT, GM, amikacin (AMK), and CPFX, qualifies as a multidrug-resistant *P. aeruginosa* (MDRP) under international criteria, exhibiting resistance to all three major antibiotic classes: carbapenems, aminoglycosides, and fluoroquinolones.

Following leukopenia induction via two injections of cyclophosphamide, mice were challenged with a clinical *P. aeruginosa* isolate and received intravenous mRNA treatment 30 minutes post-infection (**Fig. 8A**). In this immunocompromised model, all mice infected with KN41 (**Fig. 8B**) or KP2401 (**Fig. 8C**) and treated with scFv-control mRNA died within 2–3 days. In mice infected with KN41, mRNA antibody treatment significantly improved survival, with survival rates of 60% for scFv-m166 mRNA and 36.4% for scFv-m166-mFc mRNA one week post-challenge. Colistin sulfate, a curative antimicrobial agent, and mAb166 IgG protein also conferred survival benefits with 40% for colistin sulfate and 50% for m166. However, the therapeutic effects of these agents were limited in the KP2401 model, with survival rates of 18.6% for colistin sulfate, and 20% for mA166. Notably, in this challenging model, scFv-m166 mRNA achieved a survival rate of 45.5%. These results underscore the therapeutic potential of scFv antibody mRNA in a clinically relevant infection model involving multidrug-resistant highly cytotoxic strains.

**Figure 8.**
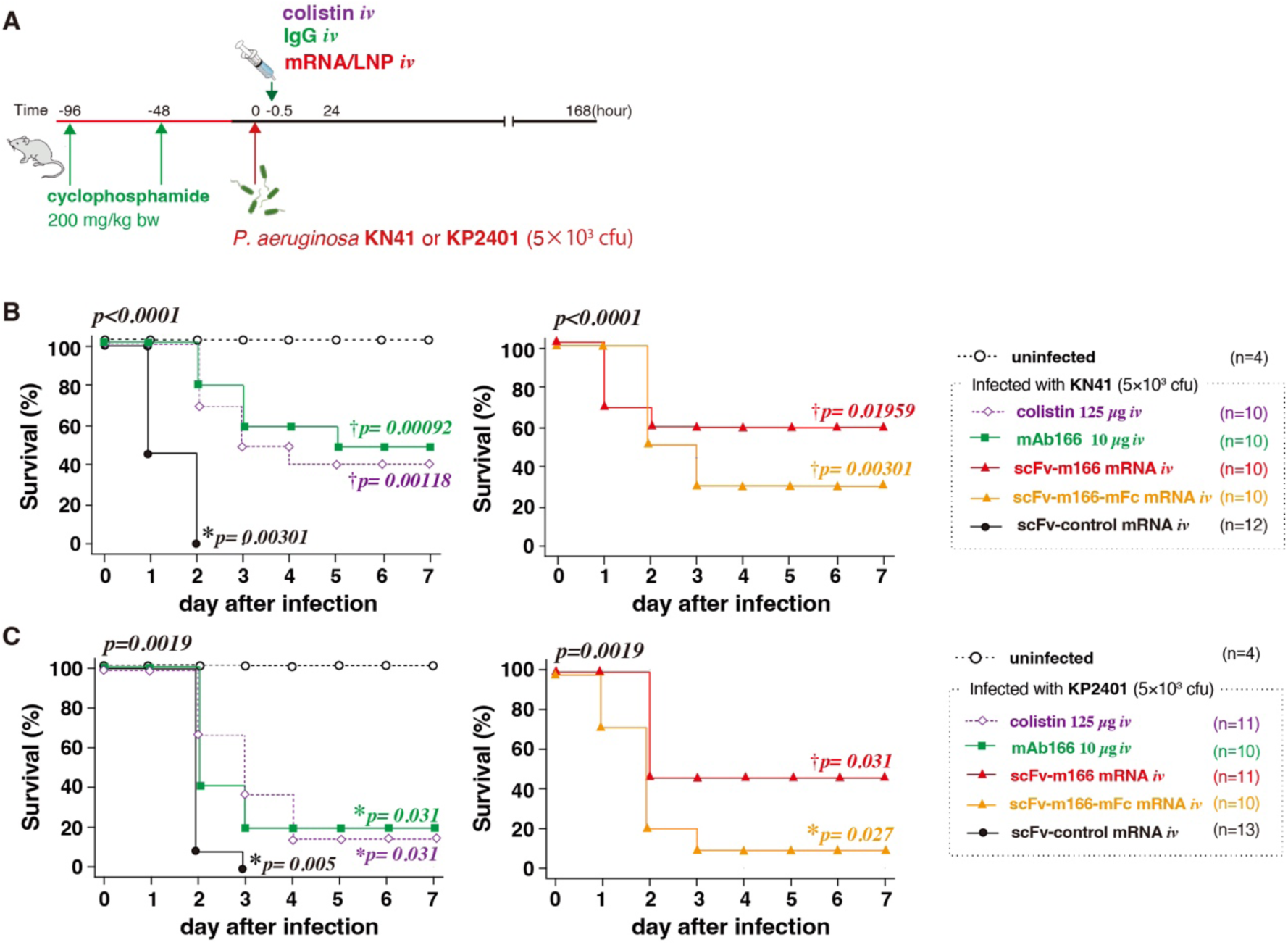
Therapeutic efficacy in immunocompromised mice infected with highly cytotoxic multidrug-resistant *P. aeruginosa*. **A.** Experimental protocol. **B.** Survival of mice challenged with a lethal dose of KN41. **C.** Survival of mice infected with KP2401.

### Spatial and temporal behavior of scFv antibodies

Upon closer observation, Fc-free scFv-m166 mRNA tended to yield higher survival rates compared to the Fc-conjugated scFv-m166-mFc mRNA in both KN41 and KP2401 (**Fig. 8**), as well as in the experiment involving 12 strains (**Fig. 7**). A similar trend was observed in therapeutic experiments using a laboratory strain, PA103 (**Fig. 4C**). In contrast, in prophylactic experiments, both Fc-free and Fc-conjugated formulations produced comparable outcomes in terms of body temperature, activity, survival rates, bacterial loads, and proinflammatory responses in the lungs (**Figs. 2, 3**). The intratracheal bacterial challenge model led to rapid infection onset, with hypothermia and reduced activity observed as early as 4 hours post-challenge (**Fig. 2C, D**). Given the acute nature of the disease, differences in the mRNA administration schedule between therapeutic and prophylactic models may critically influence treatment outcomes. Specifically, the therapeutic model may require faster antibody distribution to the infection site, the lung epithelium, than the prophylactic model. In this context, previous studies have reported that the presence of an Fc region influences antibody microdistribution within tissues^37–39^. To investigate this, we evaluated the effect of Fc conjugation on the temporal and spatial behavior of scFv antibodies in detail.

For quantifying antibodies in tissue samples using ELISA, we first generated standard curves, using the mammalian expression vectors pcDNA3.1-scFv-m166 and pcDNA3.1-scFv-m166-mFc, each incorporating a C-terminal c-Myc tag and a hexahistidine (_6×_His) tag (**Figs. S5A, S5B**). These plasmids were transiently transfected into Chinese Hamster Ovary (CHO)-K1 cells for recombinant antibody expression. The expressed scFv-m166 and scFv-m166-mFc proteins were subsequently purified in small quantities using immobilized metal affinity chromatography (IMAC) via the C-terminal His tag (**Figs. S5C, S5D**). Immunoblot analysis under non-reducing SDS-PAGE conditions confirmed that recombinant scFv-m166-mFc was produced as a dimer with an approximate molecular weight of 108.2 kDa (**Figs. S5E**). The binding affinities of scFv-m166 and scFv-m166-mFc antibodies to the PcrV antigen were assessed by ELISA, yielding half-maximal effective concentrations (EC₅₀) of 35.0 nM and 57.2 nM, respectively (**Figs. S5F**).

We conducted a time course analysis of anti-PcrV titers in the blood (**Fig. 9A**), liver (**Fig. 9B**), spleen (**Fig. 9C**), and lungs (**Fig. 9D**) of mice following intravenous injection of either scFv-m166 or scFv-m166-mFc mRNA, using ELISA on tissue homogenates. Both Fc-free and Fc-conjugated scFv antibodies became detectable in the blood and all tested organs as early as two hours post-injection. This rapid antibody production is a favorable feature of mRNA in antibody therapy, particularly for targeting acute infections. The blood and tissue levels of Fc-free scFv-m166 antibodies peaked at 3 – 4 hours post-injection. Notably, they remained detectable in the liver, spleen, and lungs for up to 24 hours. This persistence of Fc-free scFv may reflect the advantages of mRNA formulations in promoting sustained protein expression. Fc-conjugated antibodies exhibited even greater persistence, maintaining high levels even 48 hours post-injection.

**Figure 9.**
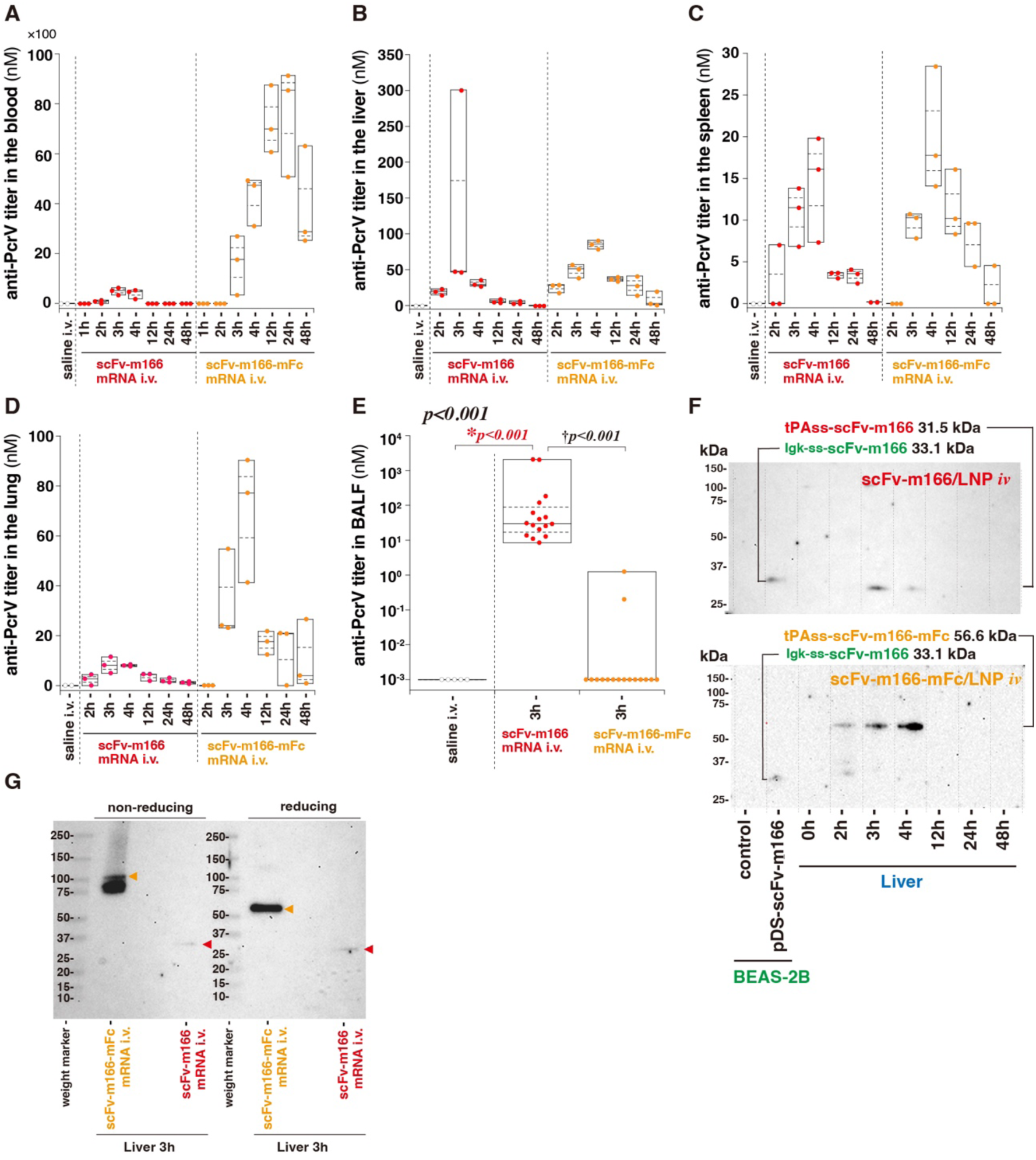
Temporal and spatial profiling of scFv antibody expression following intravenous mRNA administration. **A–E.** ELISA of anti-PcrV antibody titers, measured using plates coated with recombinant PcrV protein. **A.** Blood. **B.** liver. **C.** Spleen. **D.** Lungs. n = 3. **E.** BALF. n = 16. Box plots show the maximum, 75th percentile, median, 25th percentile, and minimum values. The lower limit of detection is 0.001 nM. The antibody titer is expressed as the equivalent concentration in nanomolar (nM), calculated based on the standard curve generated using the recombinant scFv-m166 antibody protein (**Supplementary Fig. S5**). *P* < 0.05 vs. uninfected control; †*P* < 0.05 between **scFv-m166** and **scFv-m166-mFc mRNA** groups. **F.** Time-course immunoblot analysis of scFv antibodies in liver homogenates. Cell lysate from BEAS-2B cells transfected with the eukaryotic cell expression vector pDS-scFv-m166, which expresses the scFv-m166 antibody carrying the Igκ secretion signal (Igκ-ss-scFv-m166), was used as a positive control. **G.** Immunoblot analysis of scFv antibodies in liver homogenates collected 3 hours post-injection under non-reducing and reducing conditions.

In the lungs, the primary target site for treating intratracheal *P. aeruginosa* infection, the signal from Fc-free scFv-m166 antibodies was lower than that of Fc-conjugated antibodies three hours or later post-injection. To assess antibody levels on the epithelial side, we performed bronchoalveolar lavage (BAL) three hours post-injection, as translocation of antibodies to the airway epithelium is critical for treating airway infections. Intriguingly, Fc-free scFv-m166 antibodies were significantly more abundant in BAL fluid (BALF) than Fc-conjugated antibodies (**Fig. 9E**). This result contrasts with the findings from whole lung homogenates, where Fc-conjugated antibodies showed stronger signals (**Fig. 9D**). These results indicate that the Fc-free formulation migrates from the bloodstream to the airway epithelium more efficiently than Fc-conjugated formulation.

To explain differences in tissue penetration, we evaluated the dimerization status of the antibodies in mice, using liver homogenates, as the liver showed the strongest signal among the tested organs (**Figs. 9A-D**). Time-course immunoblots of liver samples revealed strong signals 3 – 4 hours post-injection for both scFv-m166 and scFv-m166-mFc antibodies (**Fig. 9F**), consistent with the ELISA result (**Fig. 9B**). Based on these findings, we selected the 3-hour post-injection samples for SDS-PAGE under reducing and non-reducing conditions (**Fig. 9G**). Under non-reducing conditions, the size of scFv-m166-mFc antibodies was larger than under reducing conditions, while the size of Fc-free scFv-m166 antibodies remained constant between the two conditions. These results suggest that only the Fc-conjugated antibodies form a dimer in the body. Although their observed size in non-reducing conditions differed from the theoretical 108.5 kDa, dimer structure may affect electrophoretic mobility. The dimerized scFv-m166-mFc antibodies (108.5 kDa) are theoretically 3.7-fold larger than the monomeric scFv-m166 antibodies (29.2 kDa), which may explain the enhanced migration of Fc-free antibodies into the airway epithelium (**Fig. 9E**). This efficient migration might be beneficial in controlling acute *P. aeruginosa* infections (**Figs. 4, 7, and 8**).

For detailed profiling of antibody expression, we performed immunohistochemical analysis targeting the carboxyl-terminal _6×_HIS-tag of the expressed scFv antibodies (**Fig. 10**). In the liver, hepatocytes and Kupffer cells expressed scFv-m166 and scFv-m166-mFc antibodies 2-4 hours after mRNA injection (**Fig. 10A-a**). Quantitative analysis revealed significant expression in both groups (**Fig. 10A-b, c**). Splenic cells also showed positive staining three hours post-injection (**Figs. 10B, S6**). Notably, staining was observed in the cytoplasm of liver and spleen cells, presumably reflecting intracellular antibody prior to secretion. In contrast, extracellular antibodies were undetectable by this method. Lung cells did not show positive staining in either group (**Fig. 10C**), suggesting low mRNA delivery efficiency to the lungs. However, ELISA confirmed substantial antibody levels in the lungs (**Fig. 9D**), likely representing secreted antibodies. These findings suggest that mRNA LNPs induce antibody expression primarily in liver and spleen cells, followed by migration to the lungs via circulation. Importantly, these histological observations revealed no tissue damage in the liver, spleen, or lungs, supporting the safety profile of the treatment.

**Figure 10.**
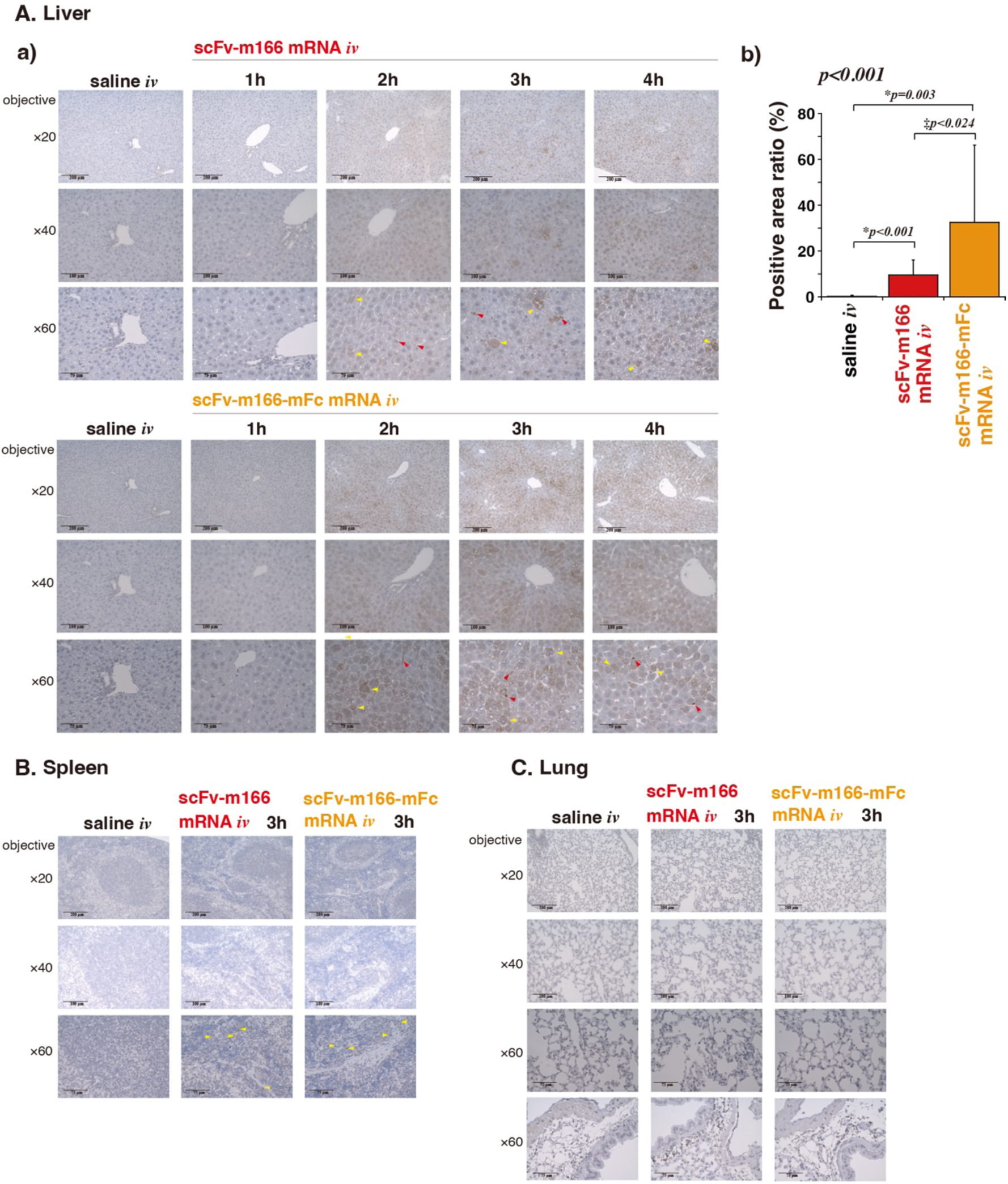
Tissue distribution of anti-PcrV scFv antibody expression following intravenous mRNA administration. Immunohistochemical staining for hexahistidine was performed to assess expression of anti-PcrV scFv following intravenous injection of scFv mRNA LNPs. **A. a)** Representative liver sections collected 1 to 4 hours post-injection. Yellow arrows indicate positively stained hepatocytes; red arrows indicate Kupffer cells. **b)** Quantitative analysis of liver sections collected 3 hours post-injection (n = 3). Quantification was performed using convolution analysis across five randomly selected microscopic fields per sample. Data are presented as mean ± SD. *P* < 0.05 vs. saline i.v. group; ‡*P* < 0.05 between scFv-m166 and scFv-m166-mFc groups. **B, C.** Immunohistochemical images of the spleen (**B**) and lungs (**C**) collected 3 hours post-injection. Yellow arrows indicate positively stained splenocytes.

To quantitatively evaluate the tissue distribution of protein expression from mRNA, we intravenously injected mRNA encoding a non-secreted form of luciferase using LNPs. Among various organs, the liver showed the highest luciferase expression, followed by the spleen (**Fig. S1A**). Other organs, including the lungs, exhibited negligible expression. This supports that mRNA LNPs primarily induce antibody expression in the liver and spleen, with antibodies subsequently migrating to the lungs via circulation. Consistent with liver histology (**Fig. 10A**), luciferase expression was observed in hepatocytes and Kupffer cells (**Fig. S1B**).

## Discussion

This study demonstrates the feasibility of antibody mRNA therapy against *P. aeruginosa* lung infection, using mRNA encoding an scFv antibody targeting the PcrV protein. Notably, our approach showed therapeutic benefits in a clinically relevant model involving immunocompromised mice infected with clinically isolated antimicrobial-resistant strains exhibiting enhanced cytotoxicity (**Figs. 7 and 8**). Clinical isolates of *P. aeruginosa* display high genetic and phenotypic diversity, reflecting the organism’s adaptability to diverse environments. Consequently, each isolate displayed distinct characteristics in terms of antimicrobial susceptibility and cytotoxicity (**Fig. 5**). In recent years, the emergence of MDRP strains resistant to all three major classes of antibiotics— carbapenems, aminoglycosides, and fluoroquinolones—has become a significant public health threat. Additionally, strains resistant to carbapenems alone are also becoming increasingly prevalent. Regarding cytotoxicity, ExoU, a T3SS-associated cytotoxin, plays a critical role by inducing cell death through disruption of eukaryotic cell membranes via its phospholipase A_2_ activity^34,40^. ExoU is a key contributor to pulmonary damage in acute lung injury^41^, and the *exoU*+ genotype is associated with bacteremia and poor clinical outcomes^42,43^. Intriguingly, anti-PcrV scFv mRNA induced significant therapeutic effects in immunocompromised mice infected with *exoU*^+^ highly cytotoxic, multidrug-resistant strains (**Figs. 7 and 8**), underscoring the potential of this approach in combating AMR.

This study also explores structural predictions of the PcrV pentamer using AlphaFold3, along with molecular interaction modeling between the PcrV pentamer and the scFv-m166 antibody (**Fig. 6**). The antibody is predicted to interact with the outlet margin at the tip of the needle structure formed by the PcrV pentamer, primarily via its VH-CDR3 domain, effectively capping the secretion pore. Our prediction aligns with recent reports demonstrating that the antibody binds to both the outer region (residues 165–223) and the inner rim of the central pore (residues 225–249) within the mushroom-shaped globular domain at the distal end of the coiled-coil shaft formed by the central and C-terminal helices of PcrV^44^. This suggests two possible mechanisms for the blocking activity of anti-PcrV antibodies: (1) inhibition of toxin extrusion through the needle by binding to the peripheral region of the pore formed by the PcrV pentamer and (2) interference with the interaction between the PcrV pentamer and the pore formed by PopB and PopD in the membrane of the target eukaryotic cell ^45,46^. The AlphaFold3-based structural interaction predictions and previous reports of amino acid substitutions in PcrV suggest that known mutations do not significantly affect the blocking activity of m166-series anti-PcrV antibodies^36^. This may explain the high therapeutic efficacy of scFv-m166 mRNA observed against clinical isolates in this study (**Figs. 7 and 8**).

Current antibody mRNA therapies generally favor Fc-conjugated formulations^20,21,23,26,31^, to prolong antigen retention via neonatal Fc receptor-mediated recycling^27^. Although a few studies have explored Fc-free scFv antibodies in mRNA-based therapies^47–50^, the comparative advantages of scFv antibodies over Fc-containing antibodies remain largely undefined. This study highlights the biodistribution advantage of Fc-free formulations. Translocation of IgG from the bloodstream to the airway epithelium is inefficient; in large animals and humans, serum concentrations of intravenously injected IgG are approximately 500-2,000-fold higher than those in BALF ^51,52^. This limits the utility of full-length antibodies in treating bacterial airway infections^53^. In contrast, Fc-free scFv antibodies, due to their smaller molecular size, can penetrate tissues more efficiently than full-length antibodies^37–39^, offering a promising solution to this distribution barrier. Indeed, in our study, the BALF concentration of Fc-free antibodies was more than 6-fold higher than that of Fc-conjugated formulations (**Fig. 9E**), despite the blood concentration of the Fc-free formulation being approximately 3-fold lower (**Fig. 9A**). This enhanced migration of Fc-free antibodies to the lung epithelium may explain their efficient therapeutic outcomes following bacterial challenge (**Figs. 4, 7, and 8**).

Notably, the benefit of Fc-free formulations in accumulating at the site of infection comes at the cost of rapid clearance from circulation, as Fc-free antibodies are typically eliminated within tens of minutes^27^. mRNA therapeutics can overcome this limitation by enabling continuous in vivo antibody production over the course of a day or more^54–56^. Indeed, Fc-free antibodies delivered via mRNA remained detectable in the liver, spleen, and lungs for up to 24 hours (**Fig. 9**). Another requirement for the Fc-free strategy is that the scFv must function independently of Fc-mediated effector functions, such as antibody-dependent phagocytosis. In this context, anti-PcrV antibodies block the transfer of type III-secreted toxins into host cells, thereby mitigating lung damage and inflammation (**Fig. 2**). Furthermore, PcrV blockade can prevent T3SS-mediated damage to alveolar macrophages, preserving their ability to clear *P. aeruginosa*^14^. These Fc-independent mechanisms likely contribute to the preventive and therapeutic effects observed in this study.

Despite the promise of our antibody mRNA therapy in controlling *P. aeruginosa*, the therapeutic advantage of scFv-m166 mRNA over mAb166 IgG protein was modest. While mRNA-based antibody therapy offers advantages in manufacturing, post-translational modification, and the use of antibody variants, such as scFvs, further improvements in therapeutic efficacy are needed. We plan to address this by fine-tuning LNPs and mRNA designs ^57–60^. Despite these limitations, this study represents a significant step forward in the treatment of *P. aeruginosa*, demonstrating successful therapy in clinically relevant mouse models using immunocompromised mice and highly cytotoxic antimicrobial-resistant clinical isolates. It also reveals the benefit of Fc-free formulations in antibody mRNA therapy. Additionally, our approach may be extended to other Gram-negative bacteria that utilize T3SS as a pathogenic mechanism ^28,29^, contributing to future strategies for combating the growing threat of AMR.

## Materials and Methods

### Anti-PcrV scFv mRNA and LNP preparation

The mRNA sequence for the single-chain anti-PcrV antibody (scFv-m166) was designed based on genetic information from our previous reports on the murine monoclonal anti-PcrV m166 IgG^9,61^. We constructed the scFv to include the essential variable regions of the heavy (H) and light (L) chains from the anti-PcrV blocking antibody m166, linked by a glycine–serine linker (Gly_4_Ser_1_)_3_. Additionally, we modified the sequence by adding a tissue plasminogen activator signal sequence (tPAss, MDAMKRGLCCVLLLCGAVFVSAR) at the N-terminus and a cMyc tag along with a 6× histidine tag at the carboxyl terminus (**Fig. 1A**). For designing the mRNA sequence of Fc-conjugated scFv (scFv-m166-mFc) antibodies, the C-terminus of tPAss-scFv was conjugated with a hinge region and CH2 and CH3 domains of mouse IgG1 followed by a cMyc tag and a 6× histidine tag downstream. Amino acid sequences are provided in **Table S2**. GenScript Japan Co. (Tokyo, Japan) prepared mRNA encoding these sequences with codon optimization, N1-pseudouridine modification, Cap1 structure, and a 100 nt poly A tail. Luciferase mRNA was purchased from Trilink Biotechnologies (San Diego, CA, USA). LNPs were formulated from ALC-0315 (MedChemExpress, Monmouth Junction, NJ, USA), ALC-0159 (MedChemExpress), 1,2-distearoyl-sn-glycero-3-phosphocholine (Fujifilm Wako, Osaka, Japan), and cholesterol (Sigma-Aldrich, St. Louis, MO, USA) at a nitrogen/phosphate ratio of 6 using microfluidics, followed by buffer exchange, according to our previous report^62^. Dynamic scattering measurement was performed using Zetasizer Nano-ZS (Malvern Instruments, Worcestershire, UK), and RiboGreen assay was performed Quant-it RiboGreen RNA Assay Kit (Thermo Fisher Scientific, Waltham, MA, USA), as described previously^63^.

### P. aeruginosa strains

In this study, a total of 12 *P. aeruginosa* strains were used, including two laboratory strains and ten drug-resistant clinical isolates. The laboratory strain PAO1 (MLST: ST549) is the reference strain for the *Pseudomonas aeruginosa* genome project and is a non-cytotoxic strain with the *exoS*⁺/*exoU*⁻ genotype^64^ ^65^. The laboratory strain PA103 (MLST: ST298) is a cytotoxic strain with the *exoS*⁻/*exoU*⁺ genotype^66^. The 12 clinical isolates were drug-resistant strains of *P. aeruginosa* obtained from hospitalized patients at Kyoto Prefectural University of Medicine Hospital (**Fig 5A**). Among them, 11 strains ̶KN5, KN15, KN17, KN27, KN28, KN33, KN34, LN37, KN41, and KP2401— were isolated between 2005 and 2014 and exhibited resistance to two classes of antibiotics^35^. Each of the four *exoS*^+^ clinical isolates had a distinct MLST. Among the six *exoU*^+^ clinical isolates, three strains—KN17, KN28, and KN41—were identified as ST235. Notably, KN28 was positive for the class 1 integron, whereas KN41 was negative. Although both KN17 and KN28 were also classified as ST357, they showed different antimicrobial susceptibility profiles: KN17 was resistant to PIPC and CAZ, while KN28 remained susceptible to both agents. The clinical isolate KP2401 (MLST: ST5295), collected in 2024, has an *exoS*⁻/*exoU*⁺ genotype, carries a class 1 integron cassette, and displays resistance to carbapenems, amikacin, and fluoroquinolones, classifying it as an MDRP strain. Bacteria from frozen stocks were streaked onto trypticase soy agar plates and cultured in trypticase soy broth supplemented with 10 mM nitrilotriacetic acid (Sigma-Aldrich) at 32°C for 13 hours in a shaking incubator. The cultures were then centrifuged at 8,500 × g for 10 minutes, and the bacterial pellet was washed twice with saline before being diluted to the desired number of colony-forming units (CFUs) per milliliter, as determined by spectrophotometry. The bacterial count was confirmed by plating diluted aliquots onto sheep blood agar and counting CFUs. The MLST types, exoenzyme genotypes, and amino acid substitutions in the *pcrV* gene of these clinical isolates were determined based on previously published reports^35^. Phylogenetic analysis based on PcrV amino acid sequences was performed using the EMBL-EBI Clustal Omega tool (https://www.ebi.ac.uk/jdispatcher/msa/clustalo), and MLST eBURST analysis was conducted using PHILOViZ 2.0 (https://www.phyloviz.net). Structural modeling of the interaction between PcrV and the antibody was carried out using the AlphaFold 3 Server (https://alphafoldserver.com) ^67^ and VMD (ver. 1.9.4a57, Biological Modeling, Carnegie Mellon University, https://biologicalmodeling.org/coronavirus/VMDTutorial), where the complex was visualized as a molecular surface using the “Surf” rendering method and “Fragment” coloring scheme. For PcrV immunoblotting, the 10 µL of supernatant from overnight bacterial culture was collected and mixed with reducing condition (2× Laemmli sample buffer containing 50 mM DTT, Invitrogen NuPAGE 10× Sample Reducing Agent, #NP0009, Thermo Fisher Scientific) with 85℃ for five minutes. Then, 10 µL sample solution (equivalent to an extraction from 0.67 mg of liver) was loaded into SDS-PAGE (NuPage Bis-Tris Mini Protein Gel 4-12%, #NP0321, Thermo Fisher Scientific), and transferred to a PVDF membrane (Invitrogen iBlot-3 Transfer Stacks, PVDF, mini 0.2 µm, #IB34002, Thermo Fisher Scientific) via a dry blot module (Invitrogen iBlot-3, #IB31001, Thermo Fisher Scientific). The membrane was blocked with 4% skim milk in PBS/T for 1 hour and then incubated with horseradish peroxidase (HRP)-conjugated anti-PcrV antibody (Creative BioLab, San Diego, CA, USA) at a 1:10,000 dilution in 4% skim milk in PBS/T overnight at 4℃. Chemiluminescence was induced via an HRP substrate (Western BLoT Quant HRP Substrate, #T7103A, Takara Bio, Kusatsu, Japan), and images were captured via an imaging device (Invitrogen iBright-CL1500 Imaging System, Thermo Fisher Scientific). The cytotoxicity of *P. aeruginosa* isolates was assessed on human bronchial epithelial cells (BEAS-2B cells immortalized with SV40, #CRL-3588, ATCC, Manassas, VA, USA). At 95% confluence, cells were co-cultured with bacteria (5×10^7^ CFU/mL) in a serum-free and antibiotic-free medium for two to four hours. Cell death was quantified using the CytoTox 96® Non-Radioactive Cytotoxicity Assay Kit (#G1780, Promega Co., Madison, WI, USA), which measures the release of lactate dehydrogenase (LDH) from the cytosol into the culture medium. LDH levels were determined by a colorimetric assay at an absorbance of 490 nm. Results are presented as the ratio of cell death at two to four hours post-infection. At 4 hours post-infection, 0.4% trypan blue staining was performed to visualize dead cells.

### Infection of mice with *P. aeruginosa* strains

All animal experiments were conducted with the approval of the Animal Care Committee of Kyoto Prefectural University of Medicine (Approval Nos. M2023-544 & M2024-535) or Institute of Science Tokyo (Approval No. A2023-191C5). Certified pathogen-free male ICR mice (7–9 weeks old, body weight 30 g) were obtained from Shimizu Laboratory Supplies Co., Ltd. (Kyoto, Japan). The mice were housed in cages with filter tops (5 mice per cage) under pathogen-free conditions. The experiments were carried out in three distinct settings: a prophylactic setting in normal mice (**Fig. 2A**), a therapeutic setting in normal mice (**Fig. 4A**), and a therapeutic setting in immunocompromised mice (**Figs. 7A and 8A**).

### Prophylactic setting in normal mice

For prophylactic interventions, the mice received either an intramuscular injection of scFv-m166 mRNA (10 µg), scFv-m166-mFc mRNA (10 µg), or luciferase mRNA (10 µg) or an intravenous injection of mAb166 IgG protein (5 µg), scFv-m166 mRNA (10 µg), scFv-m166-mFc mRNA (10 µg), scFv-control mRNA (10 µg), or luciferase mRNA (10 µg). mRNA was administered in the formulation of LNPs two hours prior to tracheal instillation of *P. aeruginosa* PA103 (1.0 × 10⁶ CFU, 50 µL). In one group, the PA103 suspension (1.0 × 10⁶ CFU, 50 µL) was pre-mixed with 10 µg of mAb166 IgG protein^32,61^, which was purchased from Creative Biolabs (New York, NY, USA). The body temperature, activity, and survival of the mice were monitored for 24 hours. The activity level was assessed as follows: Mice that moved immediately upon the start of observation and exited the circle (approximately 20 cm in diameter) within 3 seconds were given a score of 4. Those that exited within 5 seconds were scored as 3, those exiting within 10 seconds were scored as 2, and those that took longer than 10 seconds to exit were scored as 1. The mice that had died received a score of 0. Tracheal infections were induced via an endotracheal needle (modified animal feeding needle, 24 G; Popper & Sons, Inc., New Hyde Park, NY, USA) under brief anesthesia with inhaled sevoflurane (Sevofrane®; Maruishi, Osaka, Japan). Twenty-four hours after infection, the lungs were harvested, weighed, and homogenized in sterile containers with sterile water. The homogenates were serially diluted and plated on sheep blood agar to quantify the bacterial load in the lungs. The sensitivity limits of the bacteriological assays were 10 CFU/ml for blood and 100 CFU/g for lung tissue. The supernatants from the lung homogenates were stored to measure inflammatory cytokine concentrations and myeloperoxidase (MPO) activity.

### Therapeutic setting in normal mice

Mice first received an intratracheal instillation of PA103 (1.0 × 10⁶ CFU, 50 µL), and then thirty minutes later, they were intravenously injected with 100 µL solution of mAb166 IgG protein (5 µg or 10 µg), scFv-m166 mRNA (10 µg), scFv-m166-mFc mRNA (10 µg), or scFv-control (10 µg) in the LNP formulation, or saline in a control group. The survival of the mice was monitored for one week following the intervention.

### Therapeutic setting in immunocompromised mice

In the immunocompromised mouse series, leukopenia was induced by intraperitoneal injections of cyclophosphamide (200 mg/kg; Wako Pure Chemical Industries, Ltd., Tokyo, Japan) administered four and two days prior to the infection challenge ^68^. Blood samples were collected from the tail veins four days after the first cyclophosphamide injection. In the designated groups, total leukocyte and differential cell counts were determined via a hemocytometer, and mouse body weights were recorded for five days following the cyclophosphamide injections. In the first set of experiments, mice were intratracheally inoculated with either KN41 or KP2401 (5 × 10³ CFU in 50 µL). Thirty minutes later, they received an intravenous injection (100 µL) of LNP-formulated mRNA encoding either scFv-control (10 µg), scFv-m166 (10 µg), or scFv-m166-mFc (10 µg). Control groups received intravenous administration of either colistin sulfate (125 µg; 5 mg/kg) or mAb166 IgG protein (10 µg). In the second set, each of 12 clinical isolates (PAO1, PA103, KN5, KN15, KN17, KN27, KN28, KN33, KN34, LN37, KN41, or KP2401; 2.5 × 10³ CFU in 50 µL) was intratracheally administered to a pair of mice. Thirty minutes after infection, mice were intravenously treated with LNP-formulated scFv-control mRNA (10 µg) or scFv-m166 mRNA (10 µg). Survival was monitored for one week.

### Enzyme-linked immunosorbent assay quantification of anti-PcrV titers

After intravenous injection of scFv-m166 and scFv-m166-mFc mRNA in the LNP formulation, the mice were euthanized at predetermined time points, and blood, livers, spleens, and lungs were collected. The livers, spleens, and lungs were homogenized in tissue lysis solution (15 mL/g tissue; T-PER, Tissue Protein Extraction Reagent #78510, Thermo Fisher Scientific) and centrifuged to collect the supernatant for antibody titer measurement. Microwell plates (Nunc C96 Maxisorp, Thermo Fisher Scientific) were coated with rePcrV (1.0 μg/mL in 0.05 M NaHCO₃, pH 9.6) for two hours at 4°C. The plates were then washed twice with phosphate-buffered saline (PBS) containing 0.05% Tween-20 (P9416, Sigma‒Aldrich) (PBS/T) and blocked with 200 μL of 1% bovine serum albumin/PBS overnight at 4°C. The samples (100× diluted blood or 1× tissue lysis solution) were applied to the plates (100 μL/well) and incubated for two hours at 4°C. After four washes, horseradish peroxidase (HRP)-labeled anti-6×His-IgG (#HRP-66005, Proteintech, Tokyo, Japan) was added at a 1:10,000 dilution and incubated for one hour at 37°C. Following six washes, 2,2’-azino-bis(3-ethylbenzthiazoline-6-sulfonic acid) (#A3219, Sigma-Aldrich) was added to the plates, which were subsequently incubated at room temperature for 30 minutes. The reaction was stopped by the addition of 0.5 M H₂SO₄ (100 μL/well), and the optical density (OD) at 450 nm was measured using a microplate reader (MTP-880Lab, Corona Electric Co., Hitachinaka, Japan). Samples with OD values greater than 0.15 at 450 nm were considered positive.

### Bronchoalveolar lavage (BAL) fluid collection

After euthanasia with deep inhalation of sevoflurane, a tracheotomy was performed; a total of 2 mL of PBS was injected into the lungs with a syringe, and the BAL fluid was collected while vibrating with a vibration machine. The recovery rate was approximately 50–70%. Following centrifugation at 1,000 rpm for 10 min, the supernatant was used for measurement of anti-PcrV titer.

### Immunoblot assay of anti-PcrV scFv

The supernatant (100 µL, equivalent to extraction from 6.7 mg tissue extract) from the liver homogenate (15 mL T-PER /g liver) of the mice was incubated with nickel‒nitrilotriacetic acid (NTA) agarose (100 µL of 50% slurry, #30210, Qiagen, Hilden, Germany) for one hour at room temperature with constant shaking. After two washes with washing buffer, the bound protein components were eluted with elution buffer, mixed with either non-reducing condition (2× Laemmli sample buffer, Bio-Rad Laboratories, Inc., #1610737, Hercules, CA, USA) or reducing condition (2× Laemmli sample buffer containing 50 mM DTT, Invitrogen NuPAGE 10× Sample Reducing Agent, NP0009, Thermo Fisher Scientific) with 85℃ for five minutes. Then, 10 µL sample solution (equivalent to extraction from 0.67 mg of the liver) was loaded into SDS-PAGE (Miniprotean 4-15% TGX gel, 4561081, Bio-Rad Laboratories, Inc.) and transferred to a PVDF or a nitrocellulose membrane (#IB34002 and #IB33002, Thermo Fisher Scientific) via a dry blot module (Invitrogen iBlot-3, #IB31001, Thermo Fisher Scientific). After blotting on a membrane, the membrane was incubated with 8% acetic acid for 15 minutes, 3% H₂O₂ in PBS with 0.01% Tween (PBS/T) for 15 minutes, blocked with 4% skim milk in PBS/T for 1 hour and then incubated with an HRP-conjugated anti-cMyc antibody (anti-Myc-tag mAb-HRP-DirecT, mouse IgG2bk, #M192-7, MBL, Tokyo, Japan) at a 1:10,000 dilution in 4% skim milk in PBS/T overnight at 4℃.

Chemiluminescence was induced via an HRP substrate (Western BLoT Quant HRP Substrate, #T7103A, Takara Bio), and images were captured via an imaging device (Invitrogen iBright-CL1500 Imaging System, Thermo Fisher Scientific). A positive control for immunoblotting was prepared from lysates of BEAS-2B cells (human bronchial epithelium immortalized with SV40, #CRL-3588, ATCC, Manassas, VA, USA) transfected with the eukaryotic expression vector pDS-m166-cMyc-6xHis via Lipofectamine 2000 Transfection Reagent (Invitrogen #11668027, Thermo Fisher Scientific).

### Cytokines and myeloperoxidase activity

IL-6 and TNF-α concentrations in lung homogenates and plasma were quantified via an ELISA kit (BD OptEIA ELISA Sets, BD Biosciences, San Jose, CA, USA). MPO activity in the lung homogenates was measured via a biochemical method as previously described ^69^.

### Histopathological assay

A designated mouse from each group was euthanized 24 hours after infection for histological analysis. After euthanasia, the lungs were perfused with 10% buffered formalin phosphate and subsequently embedded in paraffin. Hematoxylin and eosin-stained sections were examined via light microscopy. For the immunohistochemical analysis of antibody expression, the mice were euthanized after the mRNA LNP injection. Mouse livers, spleens, and lungs were excised and fixed in 4% buffered paraformaldehyde. Paraffin sections were stained with horseradish peroxidase (HRP)-conjugated anti-_6×_HIS tag monoclonal antibody (#HRP-66005, Proteintech) and developed with 3,3’-diaminobenzidine (DAB) by Biopathology Institute Co. (Oita, Japan).

### Luciferase expression assay

Four hours after the intravenous injection of LNPs, organs were collected and homogenized in a Passive Lysis Buffer (Promega, Madison, WI, USA). Luminescence was then measured by a Lumat3 LB9508 luminometer (Berthold Technologies, Bad Wildbad, Germany). The luminescence values were standardized based on protein amount. To quantify protein expression in each liver cell type, a single-cell suspension of liver cells was obtained as described previously ^70^. The cell suspension was separated by four-minute centrifugation at 50 × g, with the precipitate used for hepatocyte isolation and the supernatant for the isolation of Kupffer cells and liver sinusoidal endothelial cells (LSECs). The precipitate underwent two additional rounds of four-minute centrifugation at 50 × g to isolate hepatocyte isolation. The supernatant was processed through the MACS cell separation system (Miltenyi, Bergisch Gladbach, Germany) to isolate Kupffer cells using Anti-F4/80 MicroBeads UltraPure and LSECs using CD146 MicroBeads UltraPure. The isolated cells were then used for the luciferase assay, performed as described above.

### Generation of recombinant anti-PcrV scFv antibody

CHO-K1 cells (JCRB9018, National Institutes of Biomedical Innovation, Health and Nutrition, Ibaraki, Japan) were cultured in CH100 serum-free medium (GMEP Cell Technologies, Kurume, Japan) supplemented with 4 mM L-glutamine. Transient transfection was performed using a mammalian expression vector, pcDNA3.1-scFv-m166, encoding a recombinant scFv-m166 antibody, and the transfection reagent TransIT-CHO (Takara Bio). After three days of continued culture, the culture supernatant was collected. The supernatant was buffer-exchanged to 20 mM phosphate buffer (pH 7.0) and concentrated approximately 100-fold using a centrifugal concentrator (#88527, Pierce Protein Concentrators PES, 10K MWCO, ThermoFisher Scientific). The concentrated sample was applied to a mini-column packed with Ni Sepharose-excel resin (#17371201, Cytiva, Uppsala, Sweden) for IMAC. After washing with wash buffer, bound proteins were eluted using 500 mM imidazole-containing elution buffer, and the eluate was dialyzed against phosphate-buffered saline (PBS) using a cup dialyzer (Slide-A-Lyzer-MINI Dialysis Device, 10K MWCO, #69570, ThermoFisher Scientific). The eluted recombinant scFv-m166 protein was analyzed by SDS-PAGE, immunoblotting, and ELISA.

### Statistical analysis

Multiple comparisons of survival curves were performed via post-hoc analysis for the log-rank test, with *p* values adjusted via the Benjamini & Hochberg method in RStudio (version 2022.7.1 Build 554, R version 4.2.1, RStudio PBC). For other statistical analyses, IBM SPSS statistical software (version 29.0.2.0, IBM Corp., Armonk, NY, USA) was used. Unpaired *t*-tests were employed to compare body temperatures between groups. Lung weight, bacterial counts, MPO activity, and cytokine concentrations in the lung homogenates were assessed via Kruskal–Wallis nonparametric tests, with multiple comparisons adjusted by the Bonferroni correction. A *P*-value of < 0.05 was considered statistically significant.

## Acknowledgments

This study is supported by Leading Advanced Projects for Medical Innovation (LEAP) (21gm0010008h0001 to S.U, T.S) from Japan Agency for Medical Research and Development (AMED), Open Innovation Platform for Industry-Academia Co-Creation (COI-NEXT) Program (JPMJPF2022 S.U.) from the Japan Science and Technology Agency (JST), Grants-in-Aid for Challenging Research (Pioneering) (23K17480 to S.U., T.S.), Grants-in-Aid for Scientific Research (A) (21H04962 to S.U.), Grants-in-Aid for Scientific Research (B) (23K24435, 22H03176 to T.S), Grants-in-Aid for Scientific Research (C) (24K12205 to M.K.) from Ministry of Education, Culture, Sports, Science and Technology, Japan (MEXT), Multilayered Stress Diseases, Science Tokyo (JPMXP1323015483 to S.U.), Nanken-Kyoten, Science Tokyo (2024-kokusai 1), and Medical Research Center Initiative for High Depth Omics, Science Tokyo. We thank Erika Mochizuki (Institute of Science Tokyo) for their technical assistance.

## Author contributions

Conceptualization: S.U., T.S.; Investigation: M.K., K.K., N.L., A.K.; Project administration: S.U., T.S.; Funding acquisition: M.K., S.U., T.S.; Supervision: S.U., T.S.; Writing—original draft: S.U., T.S.; Writing—review & editing: S.U., T.S.

---

**Figure S1.**
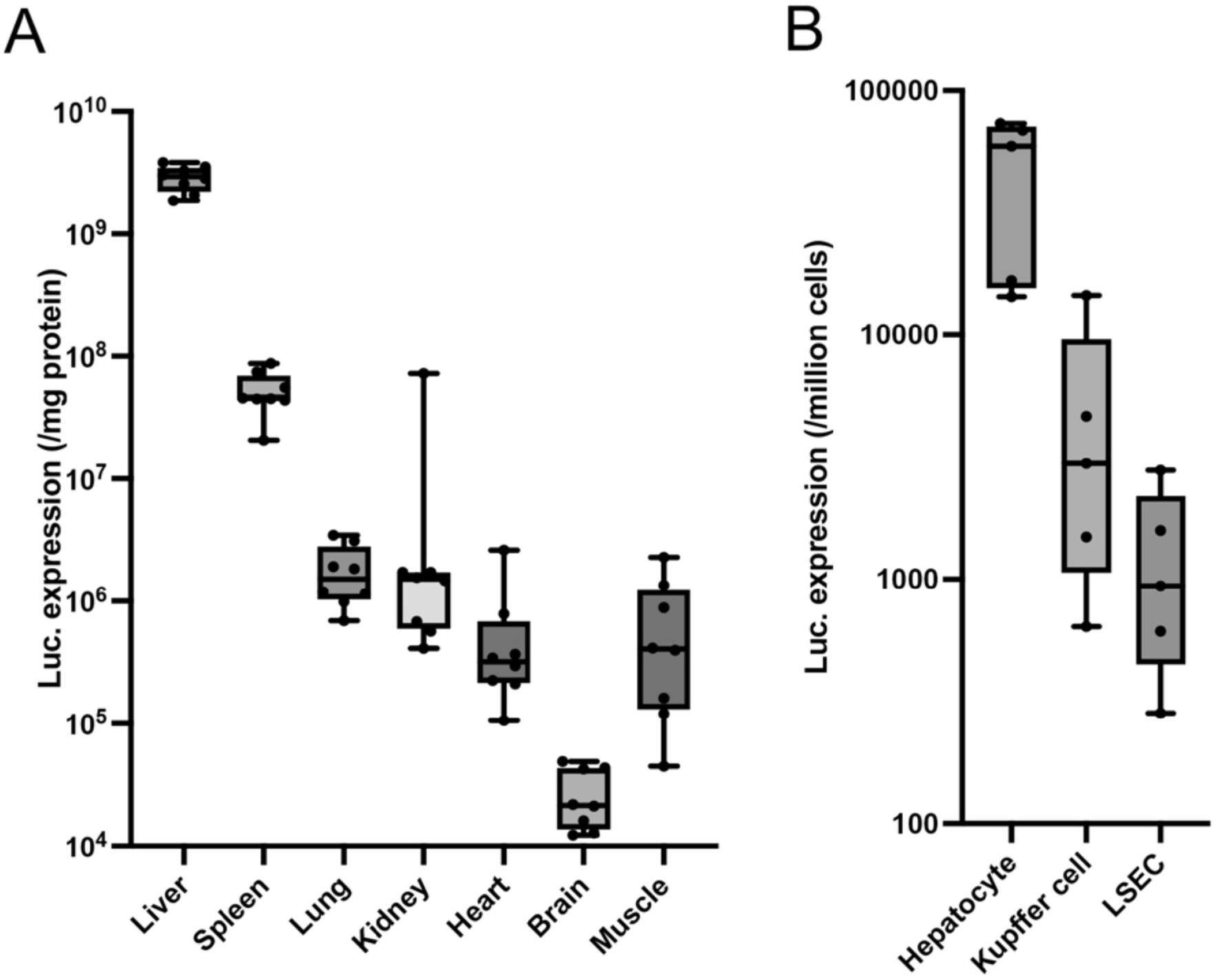
Luciferase expression following intravenous LNP injection. Expression was quantified four hours post-injection using organ homogenates (**A**) or lysates of each cell type (**B**). Box plots represent the interquartile range, the median (centerline) and the maximum and minimum (whisker lines). (**A**) *n* = 8. (**B**) *n* = 5.

**Figure S2.**
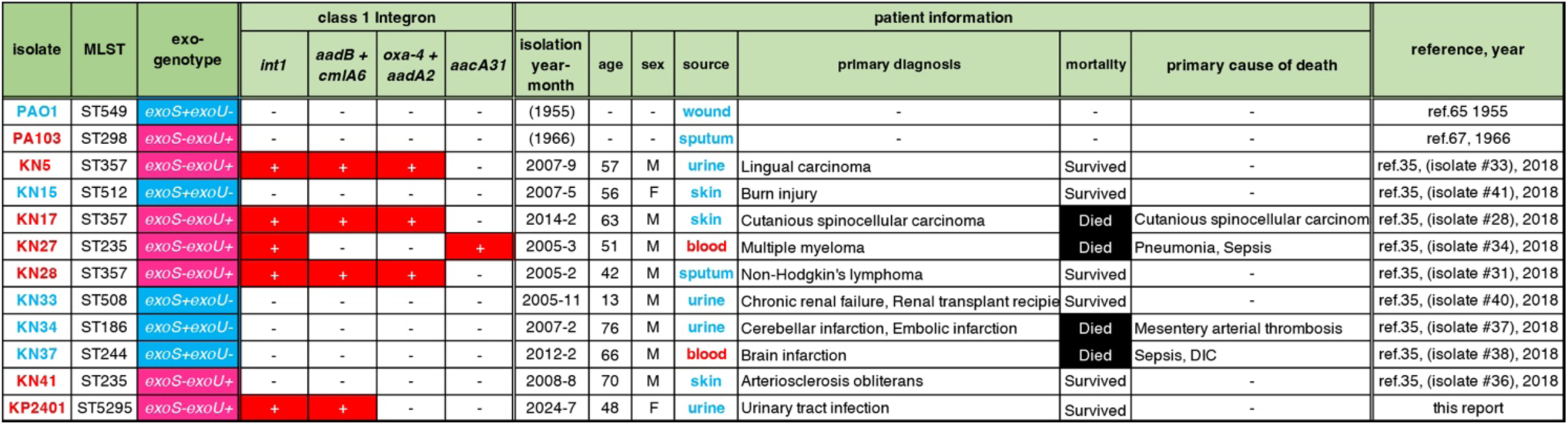
Patient information associated with the twelve isolates of *P. aeruginosa*. The PAO1 and PA103 strains are laboratory reference strains that were isolated more than 50 years ago. The eight strains from KN5 to KN41 are clinical isolates obtained at Kyoto Prefectural University of Medicine Hospital between 2005 and 2014. The KP2014 strain is a multidrug-resistant *P. aeruginosa* isolate collected at the same hospital in 2024. MLST: multilocus sequence typing, *int1*: class 1 *integron*-integrase gene, *aadB*: aminoglycoside (2”) adenylyltransferase, *cmlA6*: chloramphenicol transporter gene, *oxa-4*: β-lactamase oxallicinase-4 gene, *aadA2*: aminoglycoside (3’’) (9) adenylyltransferase, *aacA31*: AAC(6’)-31 aminoglycosides acetyltransferase.

**Figure S3.**
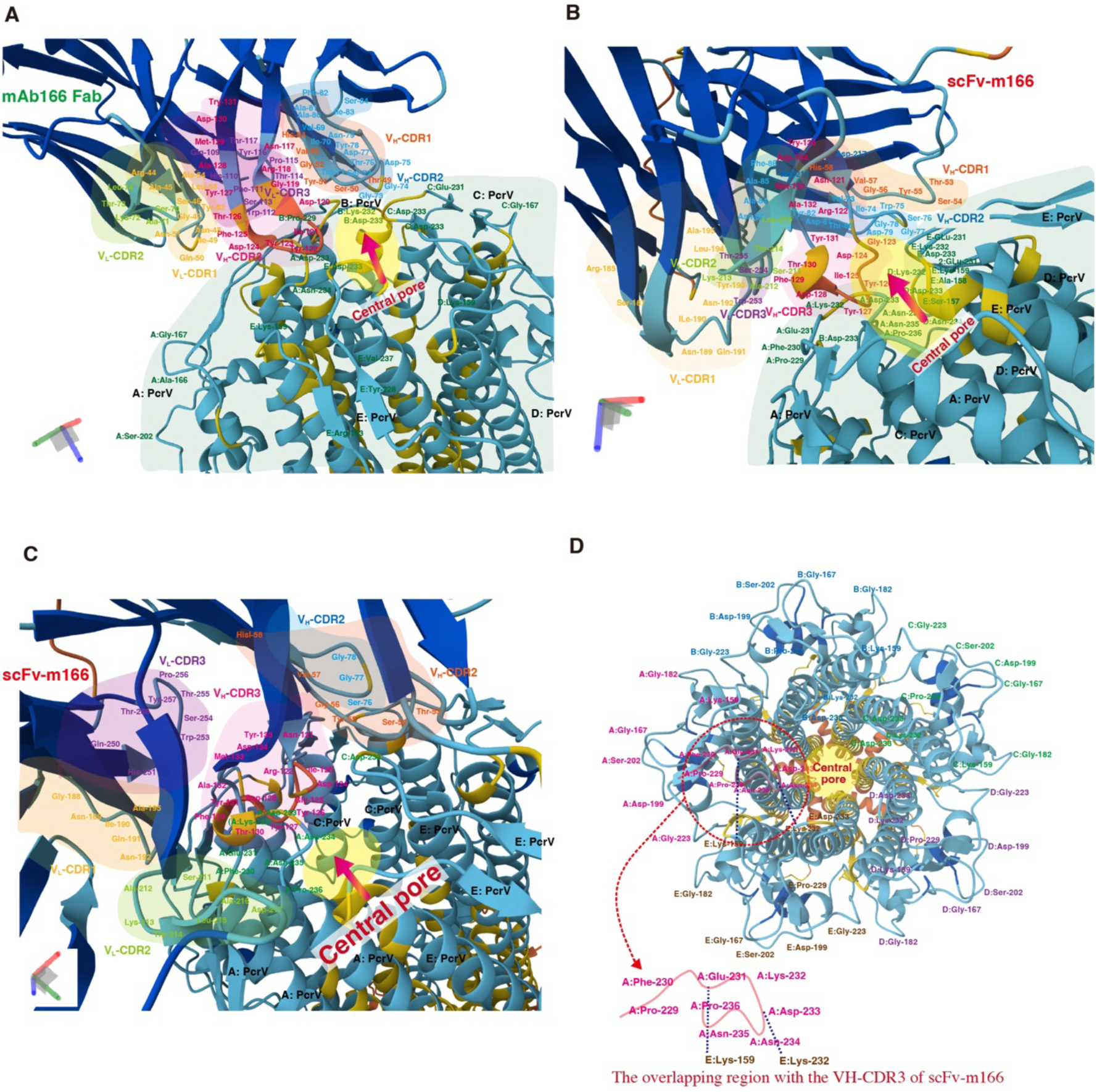
Predicted 3D-structure of pentameric PcrV binding to mAb166 Fab and scFv-m166 antibodies. Structures were predicted using AlphaFold3 multimer (DeepMind and Isomorphic Labs collaboration). **A-C.** Pentameric PcrV binding to mAb166 Fab (**A**) and scFv-m166 (**B, C**). **C.** An enlarged and slightly tilted view of (**B**). **D.** The pentameric structure of PcrV in a top view. The pore edge structure (Pro-299, Phe-230, Glu-231,Lys-232, Asp-233, Asn-234, Asn-235, and Pro-236) of PcrV overlaps with the VH-CDR3 of scFv-m166.

**Figure S4.**
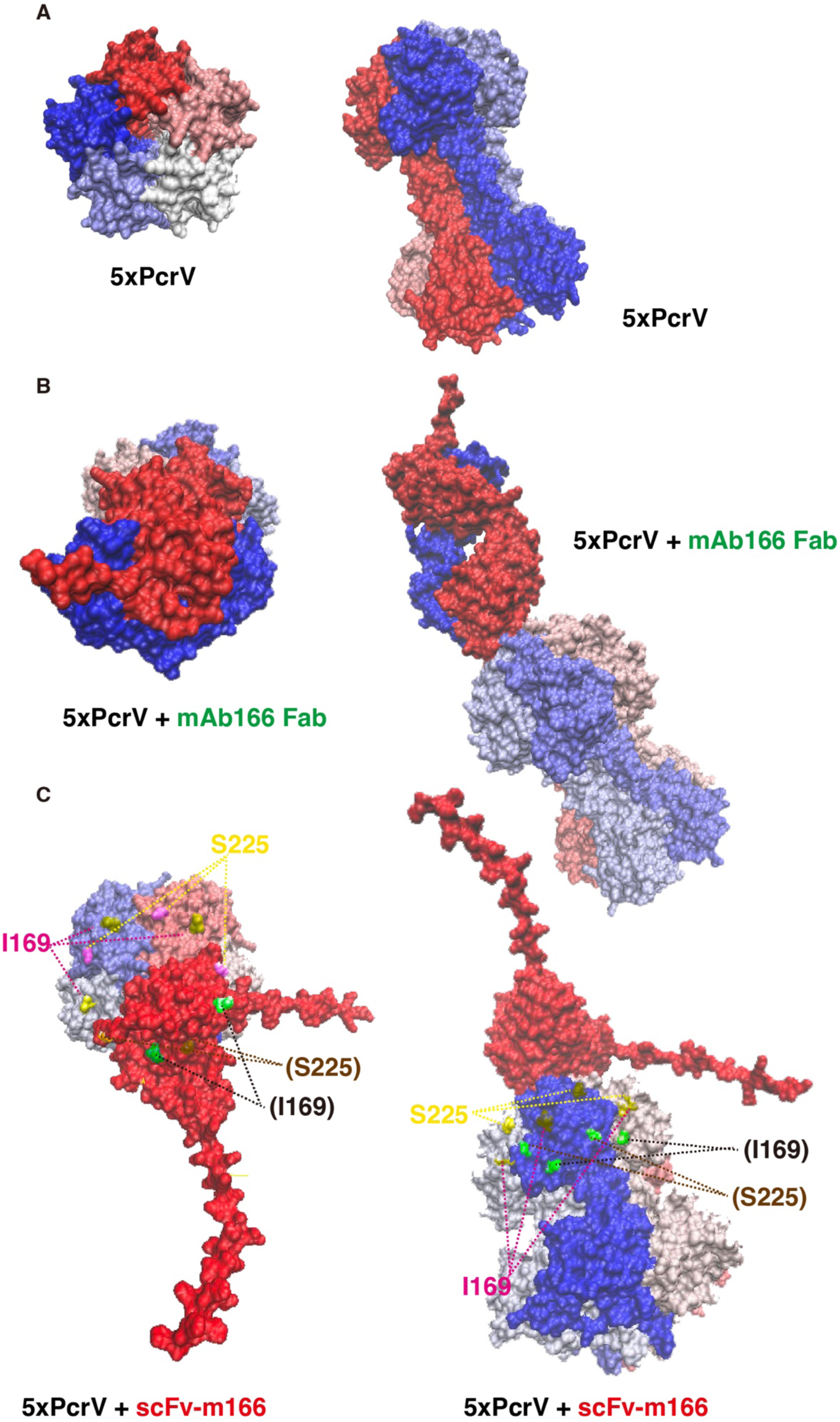
Predicted surface models of pentameric PcrV alone, with m166 Fab, and with scFv-m166 antibodies. **A.** Surface model of pentameric PcrV (left: top view; right: side view). **B.** Pentameric PcrV binding to mAb166 Fab (left: top view; right: side view). **C.** Pentameric PcrV binding to scFv-m166 antibody (left: top view; right: side view). Structural data from AlphaFold3 multimer were visualized using Visual Molecular Dynamics (VMD) where the complex was visualized as a molecular surface using the “Surf” rendering method and “Fragment” coloring scheme. Highlighted are amino acid mutations at positions #169 and #225 within the mAb166 binding epitope. The mutations shown in parentheses are located in the shadowed region of the molecular structure.

**Figure S5.**
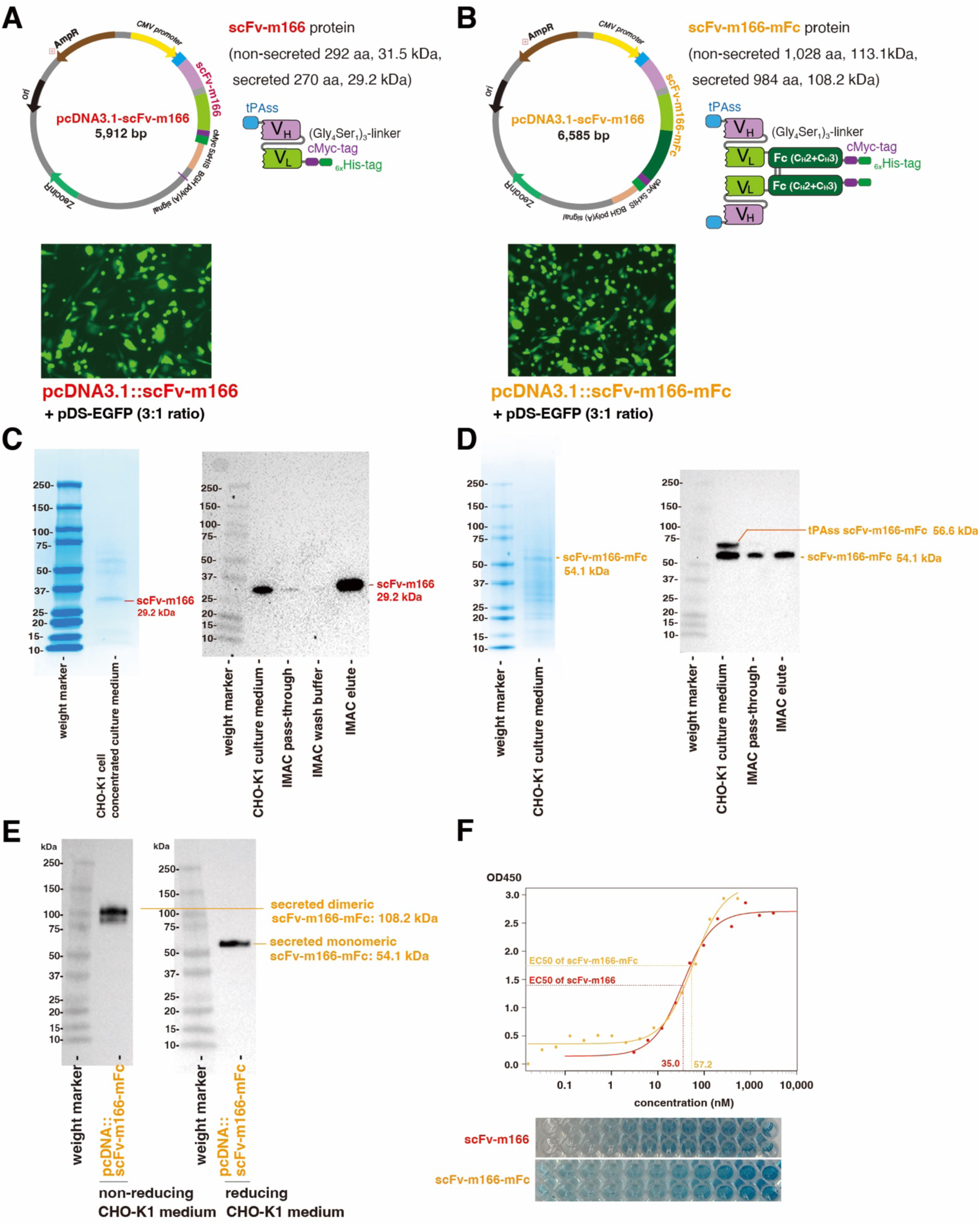
Generation of recombinant anti-PcrV scFv antibody and evaluation of its binding to the PcrV antigen. **A.** Mammalian expression vector for scFv-m166 antibody, pcDNA3.1-scFv-m166, and the structure of the expressed scFv-m166 antibody. **B.** Transfection of CHO-K1 cells with a 3:1 ratio of the mammalian expression vector pcDNA3.1-scFv-m166 and the enhanced green fluorescent protein (EGFP) expression vector pDS-EGFP. Microscopic images taken 48 hours post-transfection. **C.** (left) SDS-PAGE image of elution fractions of recombinant scFv-m166 obtained by IMAC. (right) Immunoblot analysis of recombinant scFv-m166. **D.** (left) SDS-PAGE image of elution fractions of recombinant scFv-m166-mFc obtained by IMAC. (right) Immunoblot analysis of recombinant scFv-m166-mFc. **E.** (left) Immunoblot analysis of recombinant scFv-m166-mFc after SDS-PAGE in non-reducing condition. (right) Immunoblot analysis of recombinant scFv-m166-mFc after SDS-PAGE in reducing condition. **C-D:** Detection was performed using horseradish peroxidase-conjugated anti-cMyc-tag antibody and a chemiluminescent substrate. **F.** Binding assay of recombinant scFv-m166 and scFv-m166-mFc to recombinant PcrV using ELISA. IMAC: immobilized metal affinity chromatography. ELISA: enzyme-linked immunosorbent assay.

**Figure S6.**
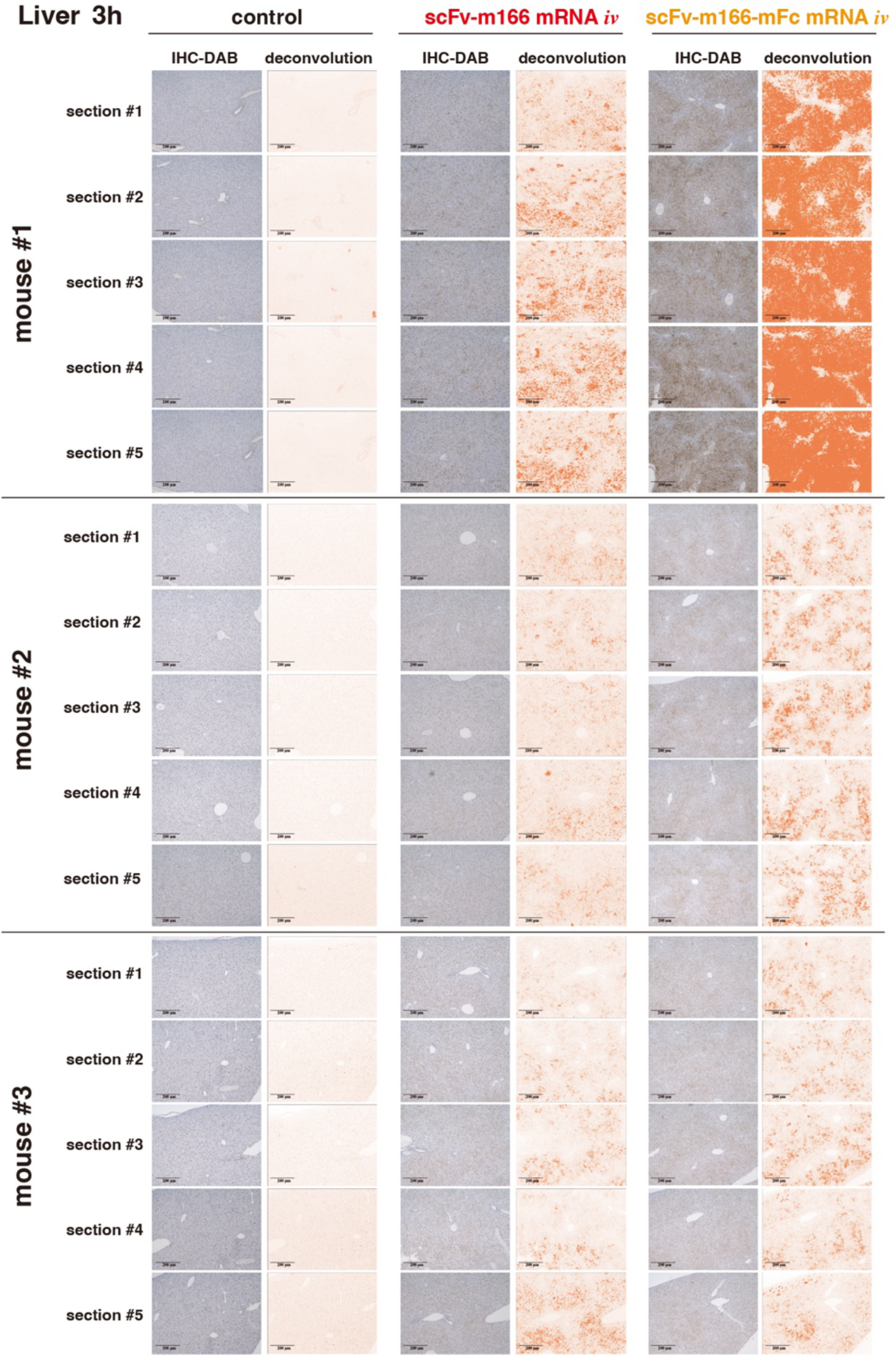
Immunohistochemical staining of anti-PcrV scFv antibody in the liver three hours post-injection. These images were used for quantification in Fig. 10A-b. DAB staining intensity was quantified using color deconvolution in ImageJ software. IHC-DAB: immunohistochemistry, 3,3’-diaminobenzidine.

**Table S1.** Characterization of LNPs.

| mRNA | Cumulant diameter<br>(nm)<sup>a)</sup> | Polydispersity index<sup>a)</sup> | Encapsulation<br>efficiency (%)<sup>b)</sup> |
| --- | --- | --- | --- |
| scFv-m166 | 124.2 ± 1.7 | 0.13 ± 0.02 | 91.5 |
| scFv-m166-muFc | 122.5 ± 0.7 | 0.12 ± 0.00 | 94.1 |
| scFv-control | 108.0 ± 0.2 | 0.14 ± 0.00 | 95.5 |

a) Determined by dynamic light scattering. Data are shown as mean ± standard error of the mean. n = 3.
b) Determined by RiboGreen assay.

**Table S2.**
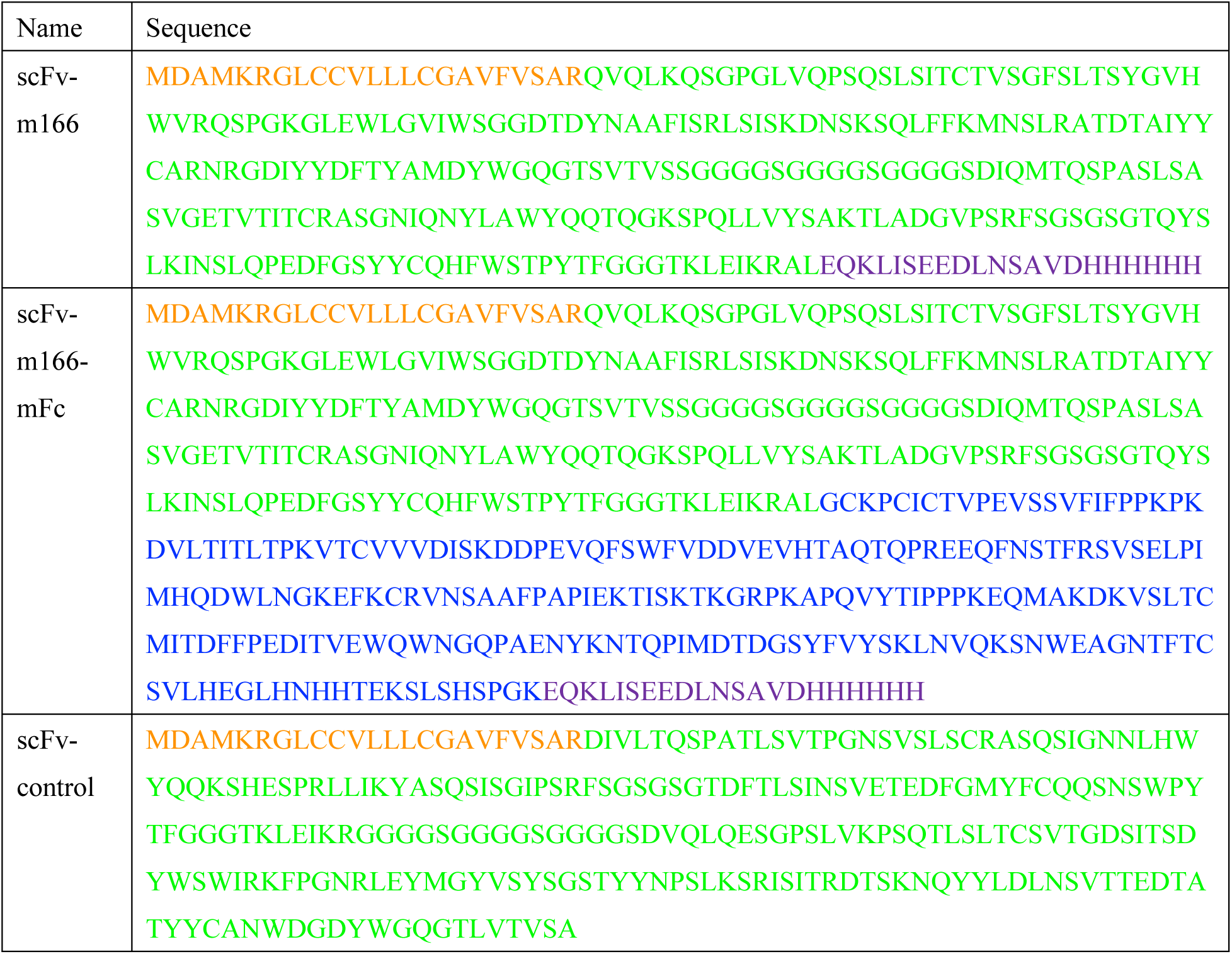
Amino acid sequences. Secretion signal (tPAss): orange; scFv: green; Fc domain (a hinge region and CH2 and CH3 domains): blue; cMyc and His tags: purple.

